# Automatic Generation of Personalised Skeletal Models of the Lower Limb from Three-Dimensional Bone Geometries

**DOI:** 10.1101/2020.06.23.162727

**Authors:** Luca Modenese, Jean-Baptiste Renault

## Abstract

The generation of personalised and patient-specific musculoskeletal models is currently a cumbersome and time-consuming task that normally requires several processing hours and trained operators. We believe that this aspect discourages the use of computational models even when appropriate data are available and personalised biomechanical analysis would be beneficial. In this paper we present a computational tool that enables the fully automatic generation of skeletal models of the lower limb from three-dimensional bone geometries, normally obtained by segmentation of medical images. This tool was evaluated against four manually created lower limb models finding remarkable agreement in the computed joint parameters, well within human operator repeatability. The coordinate systems origins were identified with maximum differences between 0.5 mm (hip joint) and 5.9 mm (subtalar joint), while the joint axes presented discrepancies between 1° (knee joint) to 11° (subtalar joint). To prove the robustness of the methodology, the models were built from four datasets including both genders, anatomies ranging from juvenile to elderly and bone geometries reconstructed from high-quality computed tomography as well as lower-quality magnetic resonance imaging scans. The entire workflow, implemented in MATLAB scripting language, executed in seconds and required no operator intervention, creating lower extremity models ready to use for kinematic and kinetic analysis or as baselines for more advanced musculoskeletal modelling approaches, of which we provide some practical examples. We auspicate that this technical advancement, together with upcoming progress in medical image segmentation techniques, will promote the use of personalised models in larger-scale studies than those hitherto undertaken.

## 1 Introduction

Musculoskeletal models have proven to be powerful computational tools to study muscle function and internal forces in healthy (Hamner et al., 2010; Saxby et al., 2016) and clinical populations (Barber et al., 2017; Fox et al., 2018; Montefiori et al., 2019b). Recent technical progress in predictive simulation approaches (Dembia et al., 2019; Falisse et al., 2019) has enabled the investigation of *”what if?”* scenarios that could support planning and execution of physical interventions. However, applications of personalised medicine often require highly accurate representations of the anatomy of the musculoskeletal system based on medical images such as magnetic resonance imaging (MRI) or computed tomography (CT) scans. For example, personalized bone geometries are essential in orthopaedics for planning surgeries and designing personalised surgical equipment (Clarke et al., 2018; Victor and Premanathan, 2013) and for creating subject-specific musculoskeletal models.

Previous studies presented methods to generate subject-specific models of the entire lower limb (Marra et al., 2015; Modenese et al., 2018) or individual joints (Barzan et al., 2019; Brito da Luz et al., 2017; Montefiori et al., 2019a; Nardini et al., 2020) and dedicated modelling tools like NMSBuilder (Valente et al., 2017a) or specialized features in the AnyBody software (Damsgaard et al., 2006) are available to implement these workflows. Nevertheless, patient-specific musculoskeletal models are currently employed in small-sized clinical applications (Falisse et al., 2020; Montefiori et al., 2019b; Pitto et al., 2019; Taddei et al., 2012; Valente et al., 2017b), mostly because the generation of each model is a time-demanding operation requiring manual intervention by specialized operators. For example, a codified approach proposed in Modenese et al. (2018) reported around 10 hours to build a complete bilateral musculoskeletal model of the lower limbs from segmented bone geometries (around two hours to create an ipsilateral skeletal model), while Scheys et al. (2006) reported on average 65 minutes to define the lower limb musculature using an atlas-based semi-automated approach. We believe that validated and fully automatic workflows are of paramount importance to enable large-scale use of these computational models.

Multiple studies with orthopaedic focus have explored the possibility of defining anatomical coordinate systems (ACSs) in the lower extremity bones based on key geometrical features. Miranda et al. (2010) and Rainbow et al. (2013) proposed automatic methods for defining ACSs for the distal femur, proximal tibia and patella, that showed minimum variability with the bone morphology. Kai et al. (2014) developed an automatic approach to identify the reference systems of the pelvis, femur and tibia based on principal axes of inertia, principal component analysis and longitudinal slicing, obtaining ACSs compatible with those created by human operators, except for the pelvis where anterior tilt was up to 18.8° higher. More recently, Renault et al. (2018) proposed multiple algorithms based on the automatic identification of the articular surfaces at the hip and knee joints, showing high repeatability of these methods when applied on 24 CT scans by three operators. However, in these previous works the ACSs were not defined consistently across publications and none of these methods has been employed for creating articulated skeletal models yet.

Statistical shape modelling workflows have recently demonstrated high potential for reconstructing bone geometries from sparse anatomical datasets (Davico et al., 2019; Nolte et al., 2016; Suwarganda et al., 2019) and landmarks digitized in the gait lab (Nolte et al., 2020; Zhang et al., 2016), but to the best of the authors’ knowledge they do not yet offer methods to generate articulated skeletal models of the complete lower limb. The bone reconstructions are limited to the long bones (Nolte et al., 2020; Nolte et al., 2016) or omit the talus and foot bones (Davico et al., 2019; Suwarganda et al., 2019; Zhang et al., 2016), and in musculoskeletal modelling contexts they have been employed to perform non-linear scaling of pre-existing muscle attachments (Nolte et al., 2016) with scarce focus towards joint modelling. Hence, a comprehensive approach to generate entire lower limb models from personalised bone geometries is still missing.

The aim of this paper is to present a tool to create models of the complete lower limb from three-dimensional bone geometries in a fully automatic way. The tool implements a sequence of operations normally performed in manual workflows, executes them in negligible computational time and generates models usable immediately in kinematic and kinetic analyses or employable as baselines for fully featured musculoskeletal models. The models produced by this tool were evaluated against manually created models employed in previous research and the joint parameters computed by competing algorithms were compared to assess their interchangeability. Differences in kinematics and kinetics curves calculated using manual and automated models were also preliminarily quantified on a set of gait simulations. Examples of further technical developments, such as joint articular mechanics and integration with anatomical models of musculature, are finally provided to demonstrate the tool’s potential for enabling large-scale studies and broader musculoskeletal research.

## 2 Materials and methods

### 2.1 Workflow to generate automatic skeletal models

A set of computational methods proposed in previous literature to define ACSs automatically (Kai et al., 2014; Miranda et al., 2010; Renault et al., 2018) were acquired and included in a more extensive modelling workflow. The geometrical methods from Renault et al. (2018) were obtained from the public “GIBOC-KNEE” MATLAB toolbox (https://github.com/renaultJB/GIBOC-Knee-Coordinate-System) and extensively modified and expanded, while the methods described by Kai et al. (2014) were independently reimplemented and those of Miranda et al. (2010) were obtained through contacting the authors of the publication. Additional algorithms were developed *ad hoc* for the purposes of this investigation (Table 1).

**Table 1.** Joint coordinate systems and available algorithms to calculate their parameters. Kai-algorithms are described in Kai et al. (2014), GIBOC-algorithms are described in Renault et al. (2018), Miranda-algorithms are described in Miranda et al. (2010) and STAPLE-algorithms are described in the Supplementary Materials of this publication. Please note that the ground coordinate system coincides with that of the medical images in all models and that the algorithms used at the pelvis were always applied to bone geometries excluding the sacrum bone (considered challenging to segment in MRI scans). The algorithms applied to the “tibia” rigid body processed the geometry of the proximal tibia, full tibia or full tibia and fibula, depending on the algorithm. Note that the “foot” rigid body, including the geometries of calcaneus and foot bones, is called “calcn” in the OpenSim models for consistency with the other models included in the software distribution.

| **Rigid Body** | **Joint Coordinate System** | **Algorithms** | **Description of joint coordinate systems used for validation (from Modenese et al. 2018)** |
| --- | --- | --- | --- |
| pelvis | ground-pelvis child | Kai-Pelvis<br>STAPLE-Pelvis | Origin: midpoint of ASIS.<br>Axes: ISB recommendations for pelvis. |
|  | hip parent | N/A | Origin: coincident with hip child origin.<br>Axes aligned with those of ground-pelvis child. |
| femur | hip child | Kai-Femur<br>GIBOC-Femur | Origin: center of femoral head (hip joint center).<br>Axes: defined as in ISB recommendations for femur. |
|  | knee parent | Kai-Femur<br>Miranda-Femur<br>GIBOC-Spheres<br>GIBOC-Ellipsoids<br>GIBOC-Cylinder | Origin: the knee joint center, as defined by selected algorithm.<br>Axes: Z axis is the medio-lateral axis as defined by the selected algorithm.<br>Y axis is perpendicular to Z, lying in the same plane of Z and the hip joint center.<br>X axis is perpendicular to Y and Z. |
| tibia | knee child | Kai-Tibia<br>Miranda-Femur<br>GIBOC-Ellipse<br>GIBOC-Centroids<br>GIBOC-Plateau | Origin: coincident with knee parent origin.<br>Axes: Z axis aligned with medio-lateral axis of knee parent<br>Y axis is perpendicular to Z lying in the same plane as talocrural-child origin<br>X axis is perpendicular to Y and Z. |
|  | talocrural parent | N/A | Origin: coincident with the talocrural child.<br>Axes: Z is aligned with the Z axis of the talocrural child.<br>Y axis is perpendicular to Z, lying in the same plane of Z and the knee joint center.<br>X axis is perpendicular to Y and Z. |
| talus | talocrural child | STAPLE-Talus | Origin: point at midpoint of the length on the axis of the cylinder fitter to the talar trochlea articular surface.<br>Axes: Z is the axis of the cylinder fitted to the talar trochlea articular surface.<br>X is perpendicular to Z, lying on a plane parallel to the foot sole XZ plane.<br>Y is perpendicular to X and Z. |
|  | subtalar parent | STAPLE-Talus | Origin: center of the sphere fitted to the articular surface of the talocalcaneal joint.<br>Axes: Z axis on the line from the center of sphere fitted to the talocalcaneal articular surface to that fitted to the talonavicular articular surface.<br>Y is perpendicular to Z, lying in the same plane of Z and the knee joint centre.<br>X axis is perpendicular to Y and Z. |
| foot | subtalar child | N/A | Origin and axes defined by subtalar parent. |
|  | foot sole (auxiliary) | STAPLE-Foot | Origin: most distal point of the calcaneus.<br>Axes: X axis pointing from the most caudal point on the talus to the midpoint of the most caudal points on the 1st and 5th metatarsal heads.<br>Y axis perpendicular to the plane identified by the points defining X.<br>Z axis is perpendicular to X and Y. |

The implemented workflow (Figure 1) consisted of the following steps: a) obtaining segmented three-dimensional bone geometries of the pelvis, femur, patella, tibia, fibula, talus, calcaneus and the other foot bones from medical images; b) automatically processing these bone models to extract the geometrical parameters required to define ACSs and appropriate joint coordinate systems (JCSs) for the parent and child bodies of each joint of the lower limb; c) creating an articulated skeletal model of the lower limb in OpenSim format (Delp et al., 2007) using the identified JCSs and ACSs.

**Figure 1.**
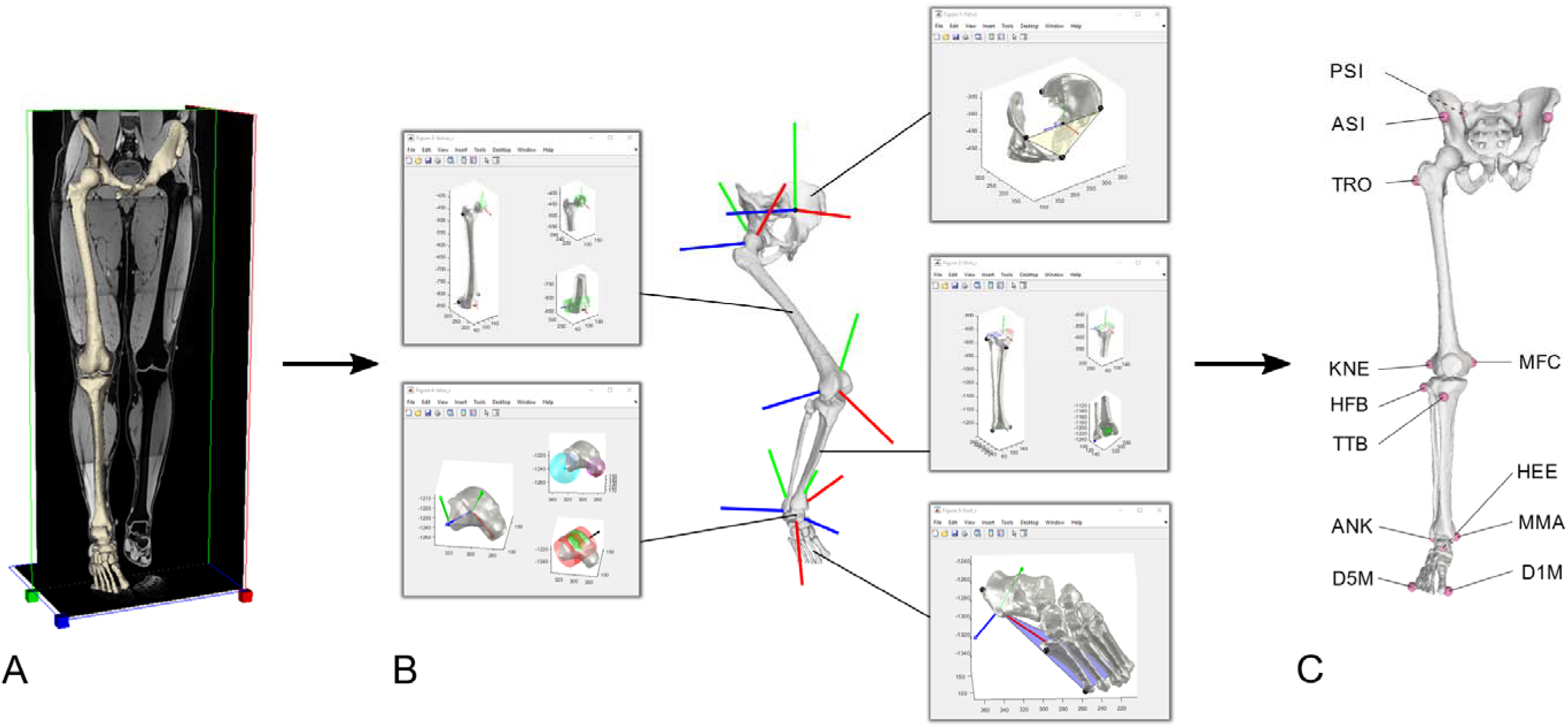
Workflow for automatic generation of articulated skeletal models: the three-dimensional bone geometries segmented from medical images (A) are fed to a MATLAB toolbox that computes the joint coordinate systems (B) used to assemble a fully functional OpenSim model inclusive of the most common bony landmarks used in gait analysis (C). For clarity only the child joint reference systems are shown in (B) for all joints.

In step b), bone geometries were analysed starting with a transformation to the ACSs defined by their principal axes of inertia (Gonzalez-Ochoa et al., 1998; Mirtich, 1996), followed by bone-specific features extraction. The complete list of algorithms available to define each JCS is reported in Table 1 and the details of the methodologies are described in their reference publications, and for the newly developed algorithms for the pelvis, talus and foot, in the supplementary materials. The articulated skeletal models were generated leveraging the MATLAB (The MathWorks, Natick, MA, USA) application programming interface (API) of OpenSim 4.1 (Seth et al., 2018): a rigid body with appropriate inertial properties (McConville et al., 1980; Winter, 2009), was created for each leg segment and joints defined based on the JCSs identified at step c). The individual JCSs, consistent with Modenese et al. (2018), are described in Table 1. For convenience in the validation step, all rigid bodies shared a local coordinate system coincident with that of the medical images, as they would have in a model generated using NMSBuilder (Valente et al., 2017a). The lower limb models included five bodies: pelvis, femur, tibia (including fibula and a rigidly attached patella), talus and foot (including calcaneus and foot bones), and five joints: a free joint between pelvis and ground (6 degrees of freedom, or “DoF”), a ball and socket joint for the hip joint (3 rotational DoF) and hinge joints for the tibiofemoral, talocrural and subtalar joints (1 rotational DoF each). No explicit patellofemoral joint was included in the models. The models also included fourteen landmarks (Table 2) automatically identified on the bone surfaces and intended for registration with the skin markers used in standard gait analysis.

**Table 2.**
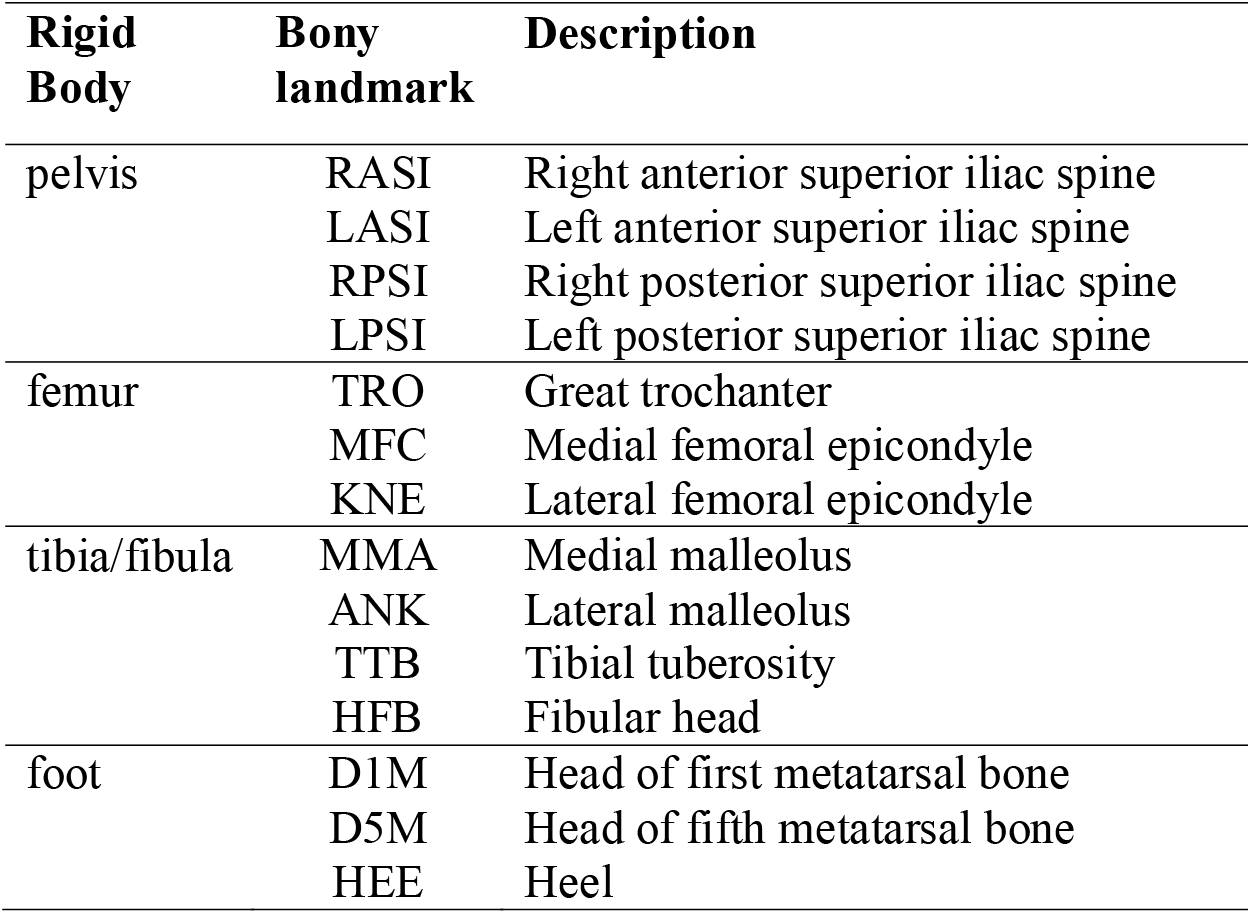
Landmarks identified on the three-dimensional bone geometries by the automated algorithms.

The entire set of scripts implementing this workflow was organized in a MATLAB toolbox named STAPLE (S*hared* T*ools for* A*utomatic* P*ersonalised* L*ower* E*xtremity modelling*).

### 2.2 Evaluation of the automatic models

The models produced using the automatic workflow were compared against musculoskeletal models of the lower limb generated for other purposes using NMSBuilder (*manual models)* and previously employed in published research (Montefiori et al., 2019b) or contributions at international conferences (Modenese et al., 2020; Modenese et al., 2019). These subject-specific models were built following the codified approach of Modenese et al. (2018) and using bone geometries available in public datasets plus an *in vivo* MRI dataset collected with the approval of the Imperial College Research Ethics Committee. A complete description of the datasets, characterized by quality of the bone geometry meshes ranging from very good (LHDL-CT) to low (JIA-MRI), is provided in Table 3 and Figure 2.

**Table 3.**
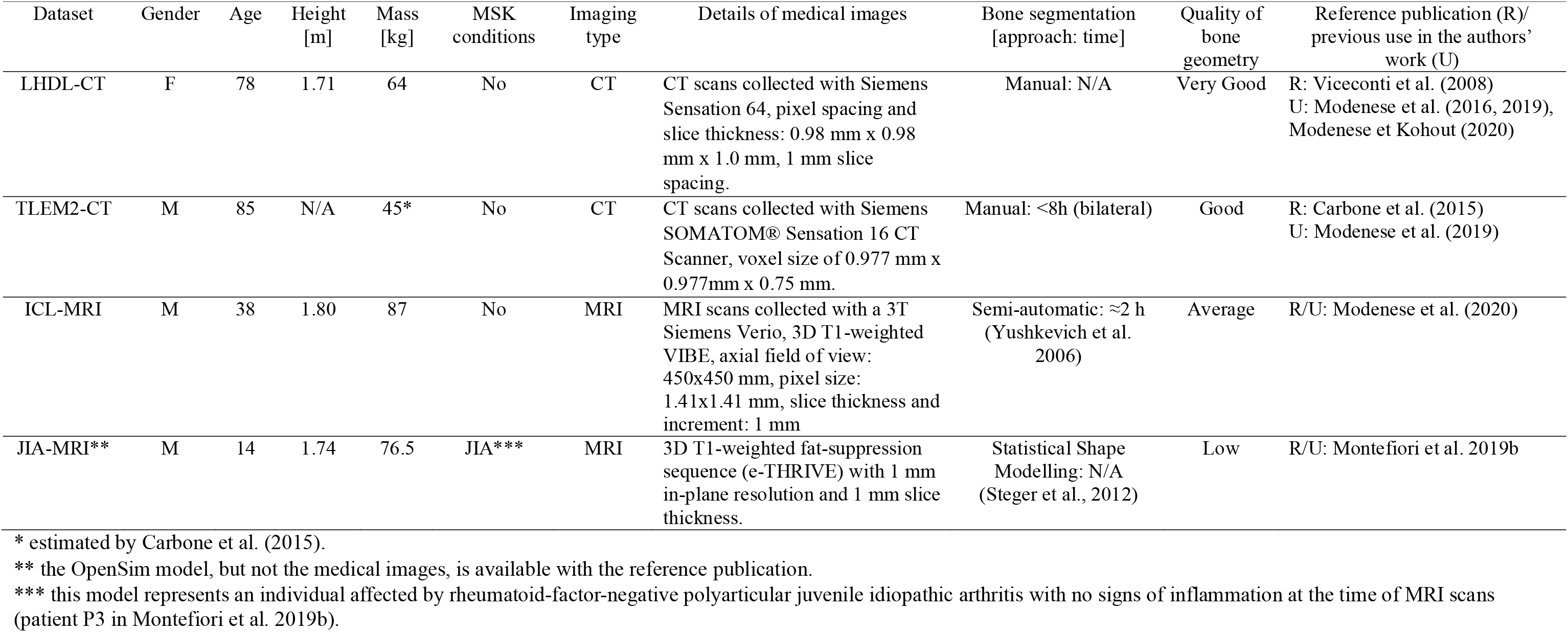
Description of the anatomical datasets employed in the current study. Please note that the LHDL-CT and ICL-MRI bone geometries were pre-processed using MeshLab. The details of the medical images are presented as reported in the reference publications. Segmentation times are reported for the datasets of which we could contact the curators.

**Figure 2.**
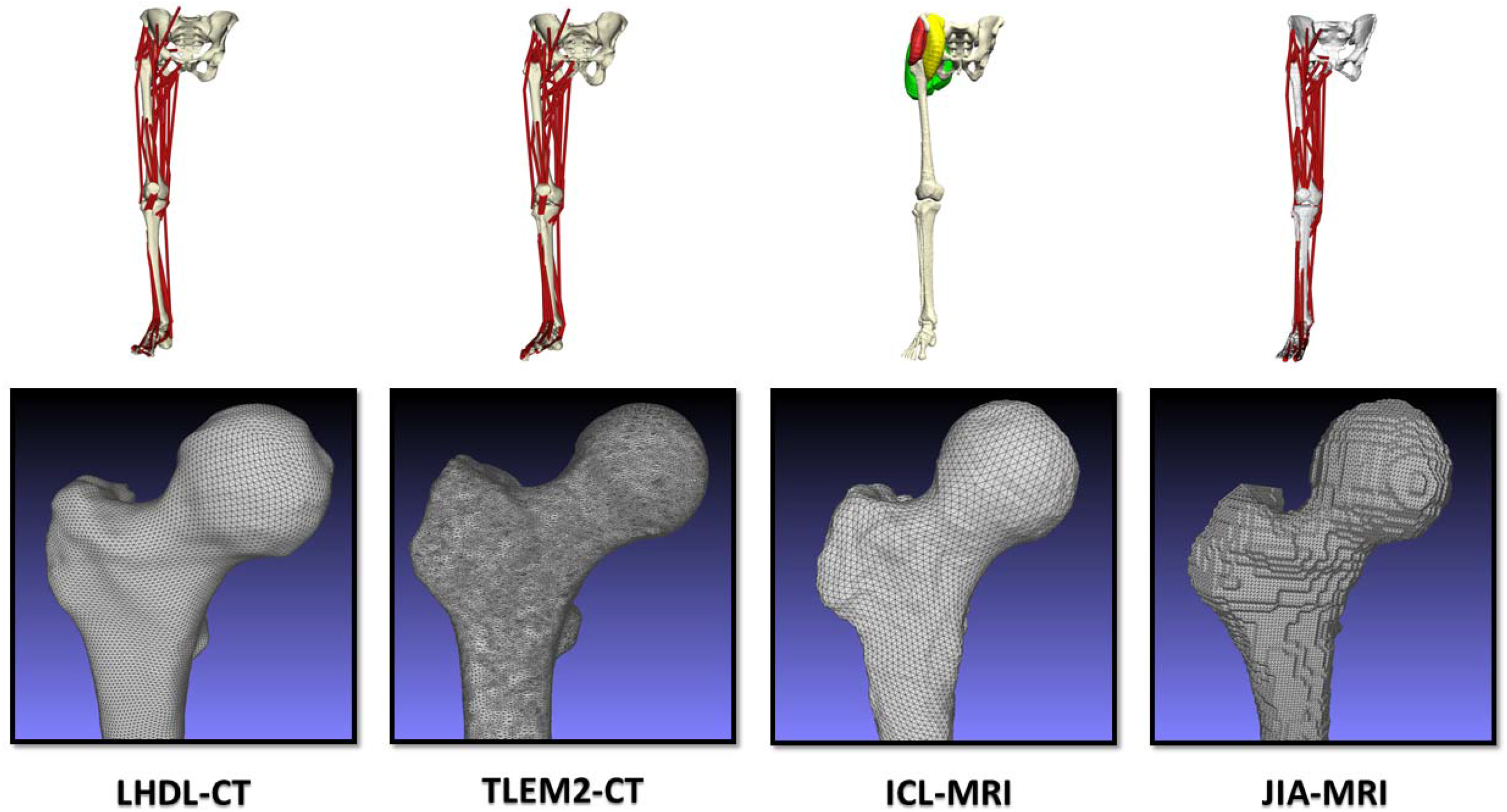
Musculoskeletal models of the lower limb from previous research used for validating the automatically generated skeletal models (first row). The models were built using bone reconstructions of variable quality (second row). Details about these models are available in Table 3.

Articulated skeletal models for all datasets were generated using the STAPLE toolbox (*automatic models*) and their JCSs compared against those of the manual models, reporting differences in their origin location and axes orientation. These quantities were evaluated in the common global coordinate system of the medical images. The automatic models were created using algorithms that matched those of the manual approach (pelvis: STAPLE-pelvis, femur: GIBOC-Cylinder, tibia: Kai-Tibia, talus: STAPLE-Talus and foot: STAPLE-Foot).

### 2.3 Comparison of joint parameters estimated by different algorithms

The JCSs of those joints for which more than one algorithm was available (ground-pelvis and knee joint) were then calculated using all the available options and the resulting JCSs compared to those employed in the evaluation part of the study, used as reference. Linear distances between origins and angular differences between axes were quantified and expressed in the JCS of the reference algorithm.

### 2.4 Gait simulations using automatic and manual models

The sensitivity of joint angles and net joint moments to JCS differences was quantified simulating six gait trials of JIA-MRI (Montefiori et al., 2019b) with the correspondent automatic and manual models, using the OpenSim inverse kinematic and inverse dynamic analyses. The resulting sets of curves were then compared using a Statistical Parametric Mapping (SPM) two-tailed t-test (significance level: α=0.05) implemented in the *spm1d* package (Pataky et al. 2013). Correlation coefficients and root mean squared errors (RMSEs) were also calculated to compare individual trials.

## 3 Results

All the automatic models employed in the study were successfully generated in less than 30s each using a standard Z640 Dell Workstation (RAM: 64 GB, CPU: 2 Intel Xeon E5-2630 2.40 GHz).

The comparison of automatic and manual models (Table 4) resulted in an overall strong similarity of the joint parameters across all considered datasets. The hip and talocrural joint centres were in excellent agreement with the manual estimations (maximum differences hip: 0.5 mm, talocrural: 1.2mm), whereas the maximum difference was 2.5 mm at the knee joint and 5.9 mm at the subtalar joint in the JIA-MRI model due to the low quality of bone reconstruction (difference range in the other datasets: 0.4-3.2 mm). The cranial-caudal position of the pelvis-ground joint origin differed by up to 4.9 mm (range: 0.8-4.9 mm) causing minor differences in pelvic tilt (range: 1.8°-3.6°, Figure 3-A) that propagated to the hip-parent JCS. The axes of the knee and talocrural hinge joints (medio-lateral Z axes) were estimated with maximum differences from manual models of 1.0° degree, while the subtalar joint axes presented maximum differences up to 2.9° in the datasets with good quality bone geometries but reached 11.3° in the JIA-MRI model (Figure 3-B).

**Table 4.** Differences between the joint coordinate systems in the manual and automatic skeletal models. The automatic models were created using the algorithms STAPLE-Pelvis, GIBOC-Femur, Kai-Tibia, STAPLE-Talus and STAPLE-Foot. Linear distances between the origins of the joint coordinate systems are expressed in the reference system of the medical images, for which axes directions are: Z pointing cranially, Y posteriorly, X to the left for LHDL-CT, TLEM2-CT and JIA-MRI; Z pointing cranially, Y anteriorly and X to the right for ICL-MRI.

| Dataset | Joint | Origin displacement vector [mm] |  |  |  | Axes Differences - Parent [deg] |  |  | Axes Differences - Child [deg] |  |  |
| --- | --- | --- | --- | --- | --- | --- | --- | --- | --- | --- | --- |
|  |  | X | Y | Z | norm | X | Y | Z | X | Y | Z |
| LHDL-CT | pelvis-ground | -0.7 | 0.8 | -3.0 | 3.1 | 0.0 | 0.0 | 0.0 | 3.1 | 3.4 | 1.2 |
|  | hip | -0.2 | -0.2 | 0.2 | 0.3 | 3.1 | 3.4 | 1.2 | 0.7 | 0.2 | 0.7 |
|  | knee | -0.8 | 1.0 | 0.2 | 1.3 | 0.7 | 0.4 | 0.7 | 0.6 | 0.4 | 0.7 |
|  | talocrural | 0.1 | 0.3 | -0.9 | 0.9 | 0.2 | 0.4 | 0.5 | 1.7 | 1.7 | 0.5 |
|  | subtalar | -0.3 | 0.3 | 0.0 | 0.4 | 0.8 | 2.6 | 2.7 | 0.8 | 2.6 | 2.7 |
| TLEM2-CT | pelvis-ground | -1.0 | 0.2 | -4.9 | 5.0 | 0.0 | 0.0 | 0.0 | 3.4 | 3.5 | 0.9 |
|  | hip | -0.1 | -0.1 | 0.2 | 0.3 | 3.4 | 3.5 | 0.9 | 0.1 | 0.2 | 0.2 |
|  | knee | 1.0 | -0.3 | 0.2 | 1.1 | 0.1 | 0.2 | 0.2 | 0.2 | 0.3 | 0.2 |
|  | talocrural | 0.1 | -0.9 | 0.4 | 1.0 | 0.7 | 0.7 | 1.0 | 1.9 | 1.6 | 1.0 |
|  | subtalar | -0.1 | 1.0 | -0.1 | 1.0 | 0.3 | 2.1 | 2.0 | 0.3 | 2.1 | 2.0 |
| ICL-MRI | pelvis-ground | -0.5 | 0.0 | 0.8 | 0.9 | 0.0 | 0.0 | 0.0 | 1.8 | 2.0 | 1.0 |
|  | hip | 0.0 | 0.1 | 0.1 | 0.1 | 1.8 | 2.0 | 1.0 | 0.1 | 2.0 | 2.0 |
|  | knee | 2.0 | 0.5 | 1.4 | 2.5 | 0.1 | 0.1 | 0.1 | 0.2 | 0.2 | 0.1 |
|  | talocrural | 0.7 | -0.6 | 0.0 | 0.9 | 0.3 | 0.4 | 0.4 | 2.5 | 2.6 | 0.4 |
|  | subtalar | -0.2 | -2.8 | -1.4 | 3.2 | 0.3 | 2.9 | 2.9 | 0.3 | 2.9 | 2.9 |
| JIA-MRI | pelvis-ground | 0.5 | 1.0 | -4.4 | 4.5 | 0.0 | 0.0 | 0.0 | 3.4 | 3.6 | 1.0 |
|  | hip | -0.2 | 0.1 | -0.4 | 0.5 | 3.4 | 3.6 | 1.0 | 0.9 | 0.1 | 0.9 |
|  | knee | 0.3 | -0.2 | -1.0 | 1.1 | 0.9 | 0.1 | 0.9 | 0.9 | 0.1 | 0.9 |
|  | talocrural | -0.2 | 0.1 | 1.2 | 1.2 | 3.0 | 3.0 | 0.2 | 2.1 | 2.1 | 0.2 |
|  | subtalar | -1.4 | -0.4 | -5.7 | 5.9 | 2.2 | 11.3 | 11.3 | 2.2 | 11.3 | 11.3 |

**Figure 3.**
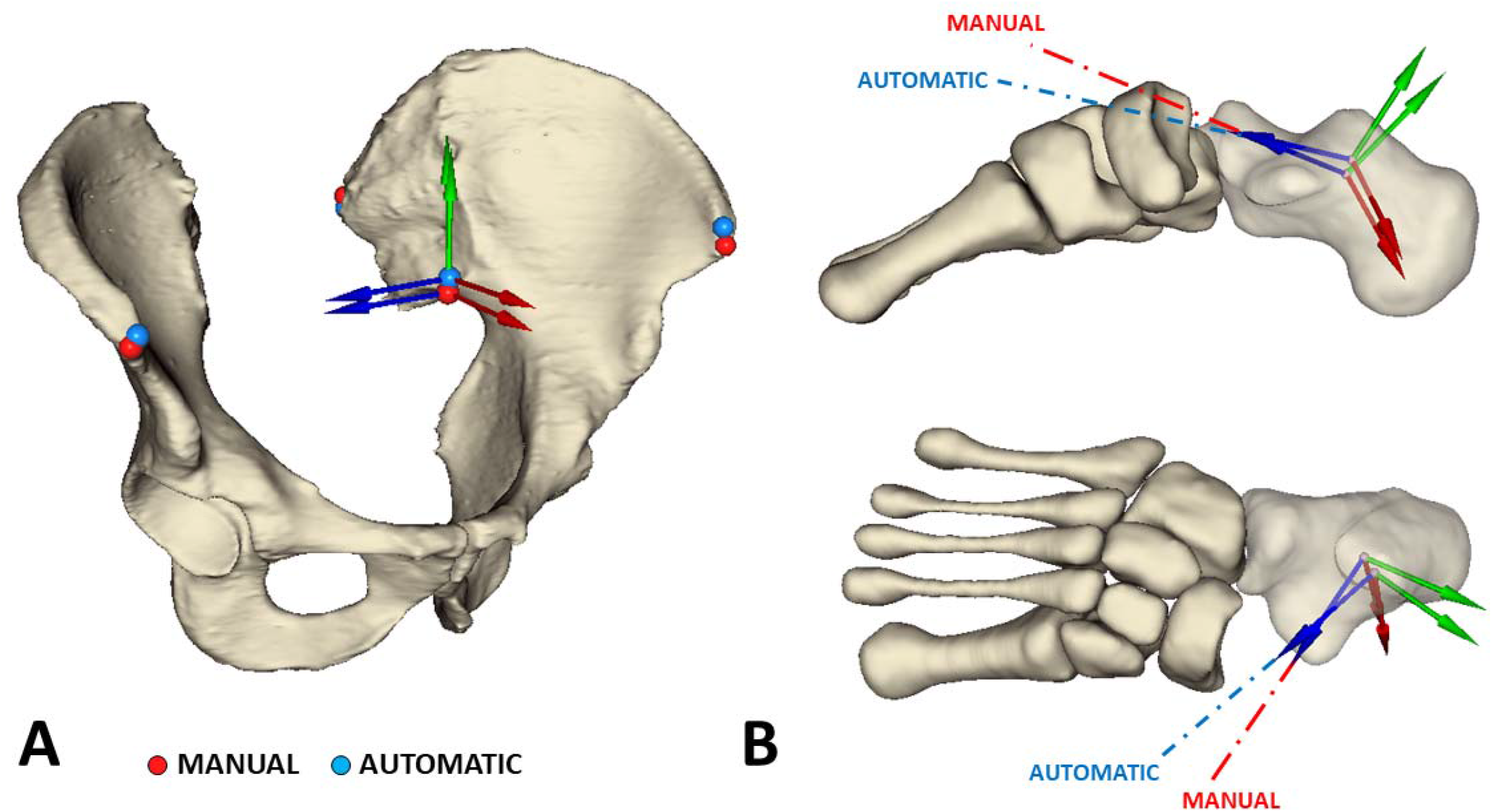
Maximum differences between manual and automatic models found at the pelvis-ground joint of TLEM2-CT using the STAPLE-Pelvis algorithm (A) and at the JIA-MRI subtalar joint using the STAPLE-Talus algorithm(B).

When comparing competing algorithms, we observed that differences among their JCSs were not always negligible (Table 5). The Kai-Pelvis algorithm presented larger pelvic tilt offsets than STAPLE-Pelvis, while GIBOC-Spheres provided the knee-parent JCSs closest to the reference algorithm (differences <1° for all cases except JIA-MRI). Kai-Femur and GIBOC-Ellipsoid resulted in more posteriorly and anteriorly located JCSs respectively, with maximum angular differences for the knee joint axis up to 4.7° (LHDL-CT) for the former and 5.2° (TLEM2-CT) for the latter. At the proximal tibia, all GIBOC algorithms computed more proximal origins than Kai-Tibia’s (range: 5.8-12.5 mm), with angular differences in the range 0.5°-11.5° for the medio-lateral axis but smaller for the proximal-distal axis (range: 0.9°-2.9°). Overall, GIBOC-Plateau and Miranda-Tibia, which failed processing TLEM2-CT, identified similar JCSs, as expected by very similar algorithms. The JCSs from GIBOC-Ellipse and GIBOC-Centroids were also found more similar to each other than to the Kai-Tibia reference.

**Table 5.** Comparison among the joint coordinate systems estimated by all the available algorithms and the specified reference algorithm. Components of the displacement vectors are expressed in the joint coordinate system of the reference algorithm. N/S means “not solved” and indicates that the algorithm crashed or computed a manifestly incorrect solution.

| Joint Coordinate System<br>[Reference Algorithm] | Algorithm | Dataset | Origin displacement vector [mm] |  |  |  | Axes Differences<br>[deg] |  |  |
| --- | --- | --- | --- | --- | --- | --- | --- | --- | --- |
|  |  |  | X | Y | Z | norm | X | Y | Z |
| ground-pelvis-child<br>[STAPLE-Pelvis] | Kai-Pelvis | LHDL-CT | -0.4 | 5.7 | 0.5 | 5.7 | 2.3 | 2.4 | 0.8 |
|  |  | TLEM2-CT | -0.2 | 1.1 | -0.2 | 1.1 | 0.5 | 0.7 | 0.5 |
|  |  | ICL-MRI | -1.3 | 5.8 | -0.1 | 6.0 | 2.1 | 2.4 | 1.3 |
|  |  | JIA-MRI | -0.1 | 0.4 | 0.1 | 0.4 | 0.2 | 0.2 | 0.2 |
| Knee-parent<br>[GIBOC-Cylinder] | Miranda-Femur* | LHDL-CT | 0.6 | -1.4 | -0.2 | 1.6 | 2.9 | 2.6 | 1.5 |
|  |  | TLEM2-CT | -3.2 | -1.7 | 0.4 | 3.7 | 0.7 | 0.9 | 0.6 |
|  |  | ICL-MRI | N/S | N/S | N/S | N/S | N/S | N/S | N/S |
|  |  | JIA-MRI | N/S | N/S | N/S | N/S | N/S | N/S | N/S |
|  | Kai-Femur | LHDL-CT | -2.4 | -0.3 | 0.9 | 2.5 | 3.6 | 3.0 | 4.7 |
|  |  | TLEM2-CT | -3.8 | -0.8 | 0.6 | 4.0 | 3.0 | 0.6 | 3.0 |
|  |  | ICL-MRI | -3.8 | -1.0 | -0.1 | 3.9 | 3.6 | 2.8 | 4.5 |
|  |  | JIA-MRI | -3.6 | -1.1 | 0.0 | 3.7 | 1.9 | 1.5 | 2.2 |
|  | GIBOC-Spheres | LHDL-CT | 0.0 | -0.5 | 1.4 | 1.5 | 0.1 | 0.1 | 0.1 |
|  |  | TLEM2-CT | 0.9 | 0.1 | 2.0 | 2.2 | 0.2 | 0.2 | 0.2 |
|  |  | ICL-MRI | 0.0 | -0.6 | 1.3 | 1.4 | 0.6 | 0.5 | 0.8 |
|  |  | JIA-MRI | -1.3 | -1.4 | 2.2 | 2.8 | 5.0 | 4.6 | 6.7 |
|  | GIBOC-Ellipsoids | LHDL-CT | 9.1 | 1.2 | 1.2 | 9.3 | 4.4 | 2.3 | 4.5 |
|  |  | TLEM2-CT | 8.6 | -0.4 | 2.0 | 8.8 | 4.4 | 3.6 | 5.2 |
|  |  | ICL-MRI | 9.0 | 0.4 | 1.6 | 9.2 | 3.6 | 1.9 | 3.6 |
|  |  | JIA-MRI | 7.8 | 1.2 | 2.1 | 8.1 | 2.6 | 3.0 | 3.4 |
| Knee-child<br>[Kai-Tibia] | Miranda-Tibia** | LHDL-CT | -0.1 | 7.5 | -1.7 | 7.7 | 10.9 | 7.0 | 9.9 |
|  |  | TLEM2-CT | N/S | N/S | N/S | N/S | N/S | N/S | N/S |
|  |  | ICL-MRI | -0.5 | 8.7 | -1.5 | 8.8 | 7.8 | 5.4 | 5.7 |
|  |  | JIA-MRI | 0.6 | 0.3 | 0.5 | 0.8 | 4.4 | 3.8 | 2.9 |
|  | GIBOC-Plateau | LHDL-CT | -1.7 | 10.4 | -1.2 | 10.6 | 9.3 | 1.7 | 9.1 |
|  |  | TLEM2-CT | -0.9 | 10.3 | -1.6 | 10.5 | 2.7 | 0.9 | 2.6 |
|  |  | ICL-MRI | -1.8 | 12.5 | -2.2 | 12.9 | 6.0 | 1.8 | 5.8 |
|  |  | JIA-MRI | -1.4 | 6.2 | -1.3 | 6.5 | 5.4 | 1.4 | 5.2 |
|  | GIBOC-Ellipse | LHDL-CT | -3.5 | 10.1 | -2.1 | 10.9 | 2.8 | 2.1 | 1.9 |
|  |  | TLEM2-CT | -2.6 | 10.0 | -2.8 | 10.7 | 4.0 | 1.2 | 3.8 |
|  |  | ICL-MRI | -4.9 | 12.1 | -2.7 | 13.3 | 2.6 | 2.3 | 1.3 |
|  |  | JIA-MRI | -4.5 | 5.8 | 0.1 | 7.4 | 6.7 | 2.0 | 6.4 |
|  | GIBOC-Centroids | LHDL-CT | -2.1 | 10.4 | -1.6 | 10.7 | 1.9 | 1.8 | 0.5 |
|  |  | TLEM2-CT | -2.4 | 10.0 | -1.5 | 10.4 | 2.3 | 1.2 | 2.0 |
|  |  | ICL-MRI | -8.7 | 11.7 | -2.8 | 14.8 | 4.2 | 2.9 | 3.2 |
|  |  | JIA-MRI | -3.3 | 6.0 | 0.0 | 6.9 | 11.7 | 1.8 | 11.5 |

\* Miranda-Femur crashed when processing JIA-MRI and fitted the shaft of the femur of ICL-MRI.
\*\*Miranda-Tibia identified inverted axes direction for TLEM2-CT and was not able to process the JIA-MRI tibial geometry unless the fibula was included as well.

The joint angles from the walking simulations performed with the manual and automatic JIA-MRI models (Figure 4) presented correlation coefficients greater than 0.99 (p<0.0001) and RMSE ≈1° degree for all coordinates except pelvis tilt and hip flex/extension, for which these errors (3.4°) were consistent with the JCS offsets in the sagittal plane (Table 4). The SPM t-test returned significant differences between the kinematics curves only for small portions of the gait cycle (from 0% to 13%), except for pelvis tilt and hip flex/extension (≈100%) and pelvic rotation (32%). The joint moments (Figure 5) had correlation coefficients larger than 0.94 (p<0.0001) and mean RMSE ranging from 0.078 (hip flex/extension) to 0.007 Nm/Kg (ankle dorsi/plantarflexion). Significant differences between the curve sets were observed mostly in the swing phase of gait, where the inertial effects were more pronounced, in clusters ranging from 11% to 37% of the gait cycle. Detailed SPM plots are available in the supplementary materials.

**Figure 4.**
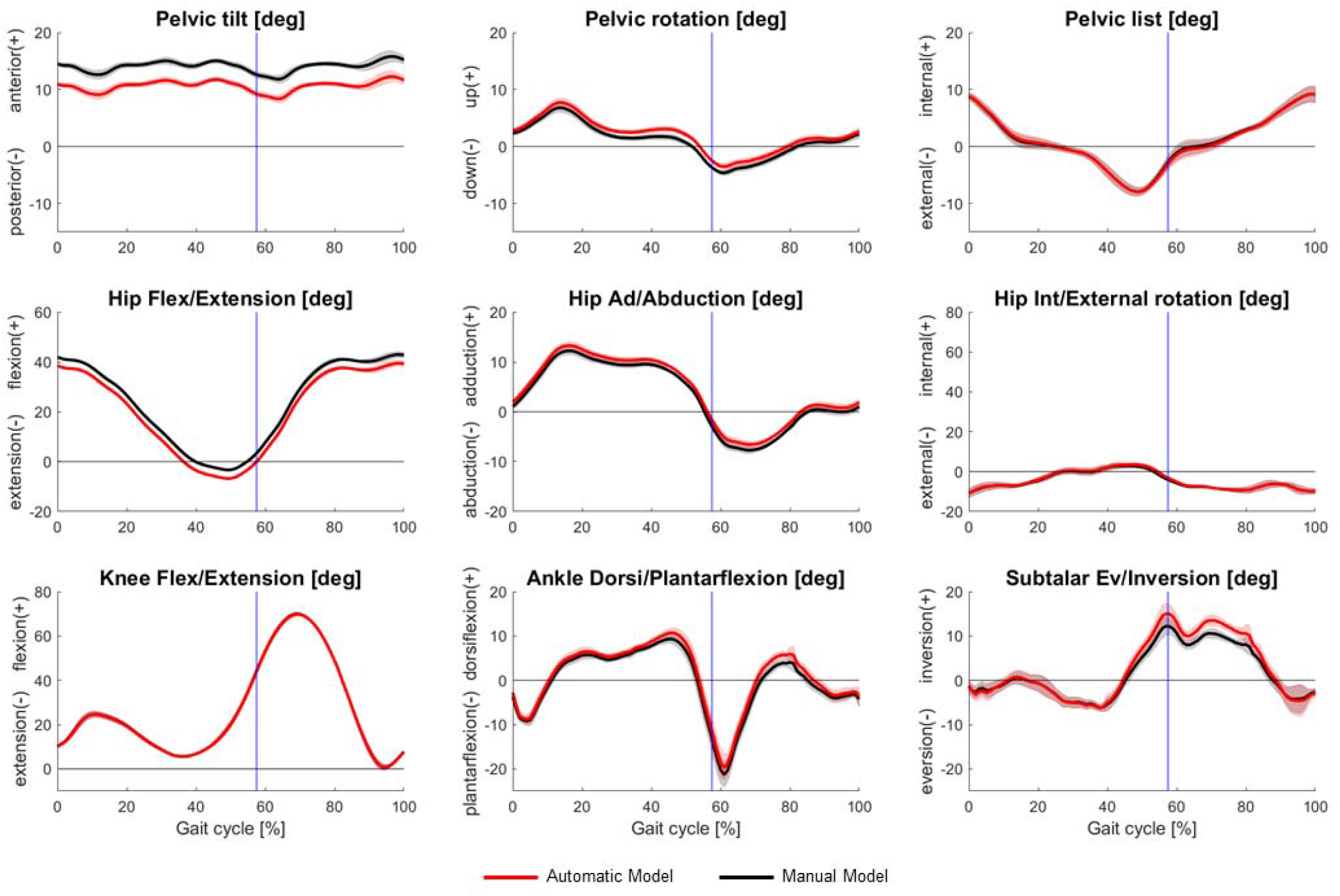
Mean joint angles (solid lines) and their standard deviations (shaded areas) calculated from six gait simulations performed with the manual (black) and the automatically created (red) JIA-MRI model. The experimental data for the walking trials were taken from supplementary materials of Montefiori et al. (2019b).

**Figure 5.**
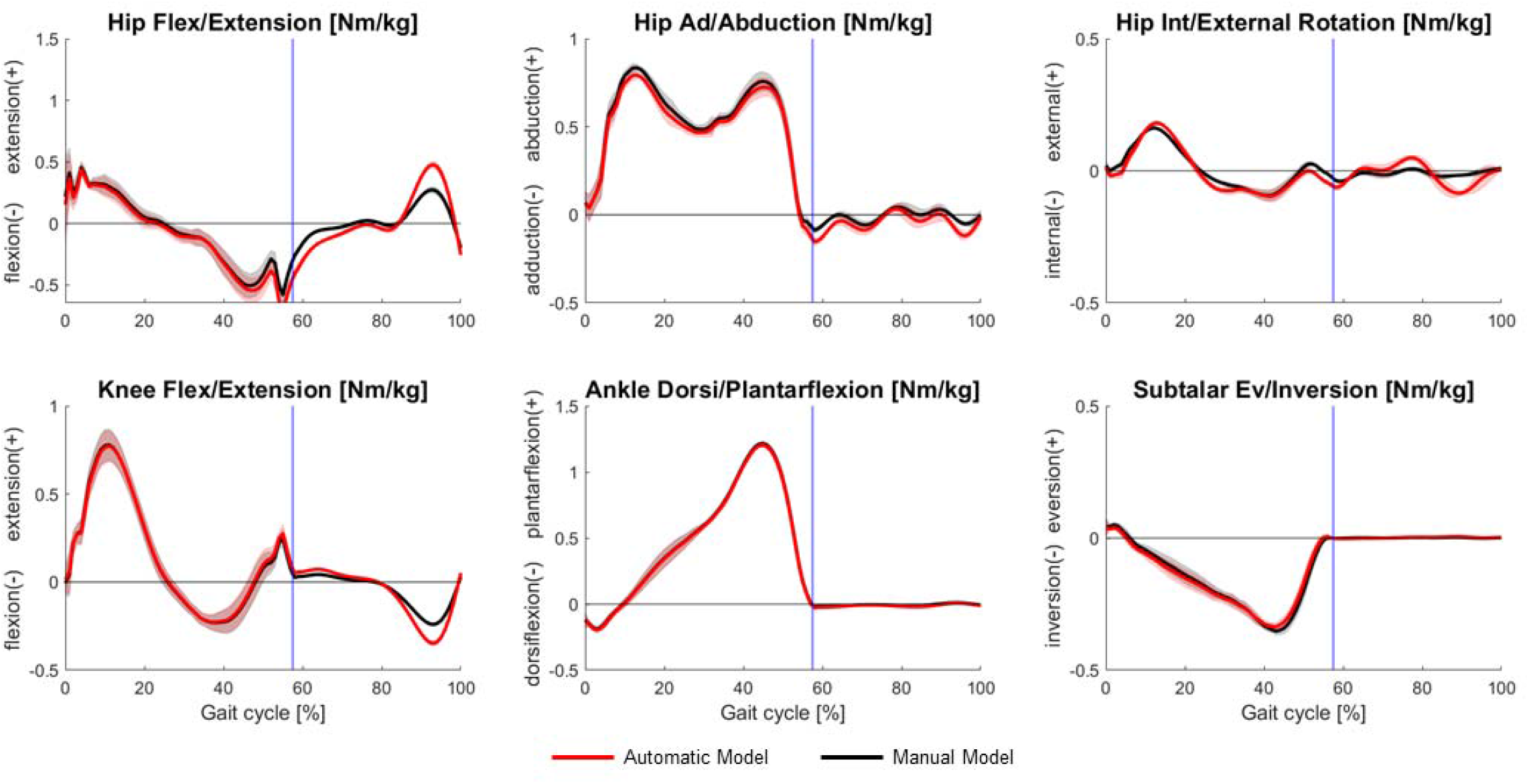
Mean net joint moments (solid lines) and their standard deviations (shaded areas) calculated from six gait simulations performed with the manual (black) and the automatically created (red) JIA-MRI model. The moments are normalised by body mass. The experimental data for the walking trials were taken from supplementary materials of Montefiori et al. (2019b).

## 4 Discussion

The aim of this paper was to present a tool to automatically create personalised models of the lower limb from three-dimensional bone geometries segmented in medical images. To evaluate the proposed methodology, we generated four automatic models and compared them against models manually created from the same data in other research projects. We found that the automatic and manual models were remarkably similar (Table 4), with largest differences observed for the pelvis-ground and subtalar joints. In the pelvis, the JCS origin was misplaced by up to 4.9 mm cranially due to the identified bony landmarks (Figure 3-A). This difference causes a systematic anterior tilt offset in the range 1.8°-3.6° that propagates to the pelvis and hip joint kinematics, as confirmed by the gait simulations. Although not negligible, this offset represents a substantial improvement (almost 15°) compared to the results of Kai et al. (2014). More recent algorithms, e.g. Fischer et al. (2019), could be considered for future additional comparisons. The automatic subtalar joint axis in the JIA-MRI dataset was also noticeably different from the equivalent manual model (11.3°, Figure 3-B). This discrepancy was attributed to the low-quality talus bone reconstruction since in all the other models the same axis was estimated within 2.9°.

JCS differences at the hip, knee and talocrural joint were well within the ranges of human inter-operator repeatability reported as standard deviations in previous investigations (Hannah et al., 2017; Martelli et al., 2014; Montefiori et al., 2019b) and even the largest difference observed at the subtalar axis was close to the maximum inter-operator variability (9.6°) reported by Montefiori et al. (2019a). These JCSs differences caused joint angles offsets in the gait simulations, as expected based on previous studies (Kainz et al., 2016; Martelli et al., 2014; Montefiori et al., 2019b; Valente et al., 2014). The largest difference in net joint moments (Figure 5) occurred for the hip flex/extension moment in the swing phase of gait and was attributed to dissimilar shank mass between models (25% difference), based on the results of Wesseling et al. (2014). This discrepancy was due to the inertial properties being estimated using segmented soft tissues geometries (not publicly available) in the manual JIA-MRI model (Montefiori et al., 2019b) and using regression equations (McConville et al., 1980; Winter, 2009) in the automatic model. It is worth highlighting that simulation results from the STAPLE models will have zero inter- and intra-operator variability due to model construction.

In all considered models, the JCSs were created as in Modenese et al. (2018), but other algorithms were also assessed (Table 1) to encourage researchers to implement different modelling workflows. At the pelvis, using Kai-Pelvis instead of STAPLE-Pelvis resulted in larger pelvis tilt offsets. For the long bones, JCSs estimated by different algorithms exhibited variability comparable with Renault et al. (2018). At the distal femur the differences among JCSs were small (<5° in the 93% of estimations) but not negligible, therefore the choice of the algorithm, e.g. fitting ellipsoids (Sholukha et al., 2011) or spheres (Yin et al., 2015) to the femoral condyles articular surface, must be justified with careful functional anatomy considerations relevant to the research question. At the proximal tibia, the mechanical axis (Y axis) was similar between Kai-Tibia and GIBOC algorithms (range: 0.9°-2.9°) but less for Miranda-Tibia (up to 7°). Larger differences were observed in the medio-lateral axis (range: 0.5°-11.5°) and anterior-posterior axis (range: 1.9°-11.7°). The workflow of Modenese et al. (2018) connects the tibia to the tibiofemoral axis identified in the femur in the segmented pose, hence when the knee joint assumes the neutral position with zero flexion/extension angle, offsets due to standard supine imaging (Hirschmann et al., 2015) could still be present. The automatic algorithms could inform procedures to adjust the tibiofemoral alignment, but further work is required in this respect.

A current limitation of the study is that our validation datasets were only four, and despite their heterogeneity, did not include bone geometries presenting abnormalities, deformities and partial geometries due to imaging (Henckel et al., 2006) or amputations that could be encountered clinically. This limitation is mitigated by the availability of multiple algorithms for the long bones, some of which specifically developed for incomplete geometries (Miranda et al., 2010; Renault et al., 2018), but additional validation is undoubtedly required. Also, the quality of the bone geometries can vary depending on the medical image modality and their specifications and resolution, potentially affecting the automatic algorithms. This possibility was not systematically investigated in this study, although we can report that, when processing low quality bone models, algorithms based on bone global geometrical features (Kai-algorithms) were generally more robust than those relying on the identification of the articular surfaces (GIBOC-algorithms), while the Miranda-algorithms seemed the most affected and failed to process some of the datasets.

The importance of a fully automatic approach as a step towards clinical application of modelling can be fully appreciated considering the reduction in total processing time for generating a skeletal model, from segmentation of the medical images to the final OpenSim model. For the ipsilateral models considered in this study, the bone segmentation and manual modelling steps required a comparable time (between two and four hours each, depending on the experience of the operator). For example, an expert operator segmented the ICL-MRI dataset, the only one that we processed directly, in around two hours using the semi-automatic functionalities of ITK-Snap (Yushkevich et al., 2006) and created an ipsilateral skeletal model in roughly the same amount of time. Based on the TLEM2 segmentation time (Table 3), our previous experience (Renault et al., 2018) and literature reports (Matsiushevich et al., 2019), we estimated similar processing times also for CT datasets. STAPLE executes modelling workflows in seconds, therefore practically halving the total processing time to generate a skeletal model and reducing it essentially to an image segmentation task. This means, for example, that when radiological scans are collected and segmented for planning musculoskeletal surgical interventions, obtaining further model-based biomechanical analyses becomes a quick and inexpensive option. Considering that deep learning techniques are reducing also the segmentation time by order of magnitudes, e.g. Noguchi et al. (2020) reported ~12s for segmenting a full-body CT scan of ~600 slices, the generation of personalised lower limb models in a number of clinical applications appears technically feasible.

The STAPLE toolbox is intentionally modular and can automate the generation of entire or partial lower limb models (see supplementary materials for examples). Moreover, the GIBOC and STAPLE algorithms provide a large amount of anatomical information such as articular surfaces (Figure 6-A) and bone profiles that can be used for implementing more advanced joint models than those proposed here, e.g. contact models (Brandon et al., 2017; Conconi et al., 2015) or parallel mechanisms (Sancisi and Parenti-Castelli, 2011).

**Figure 6.**
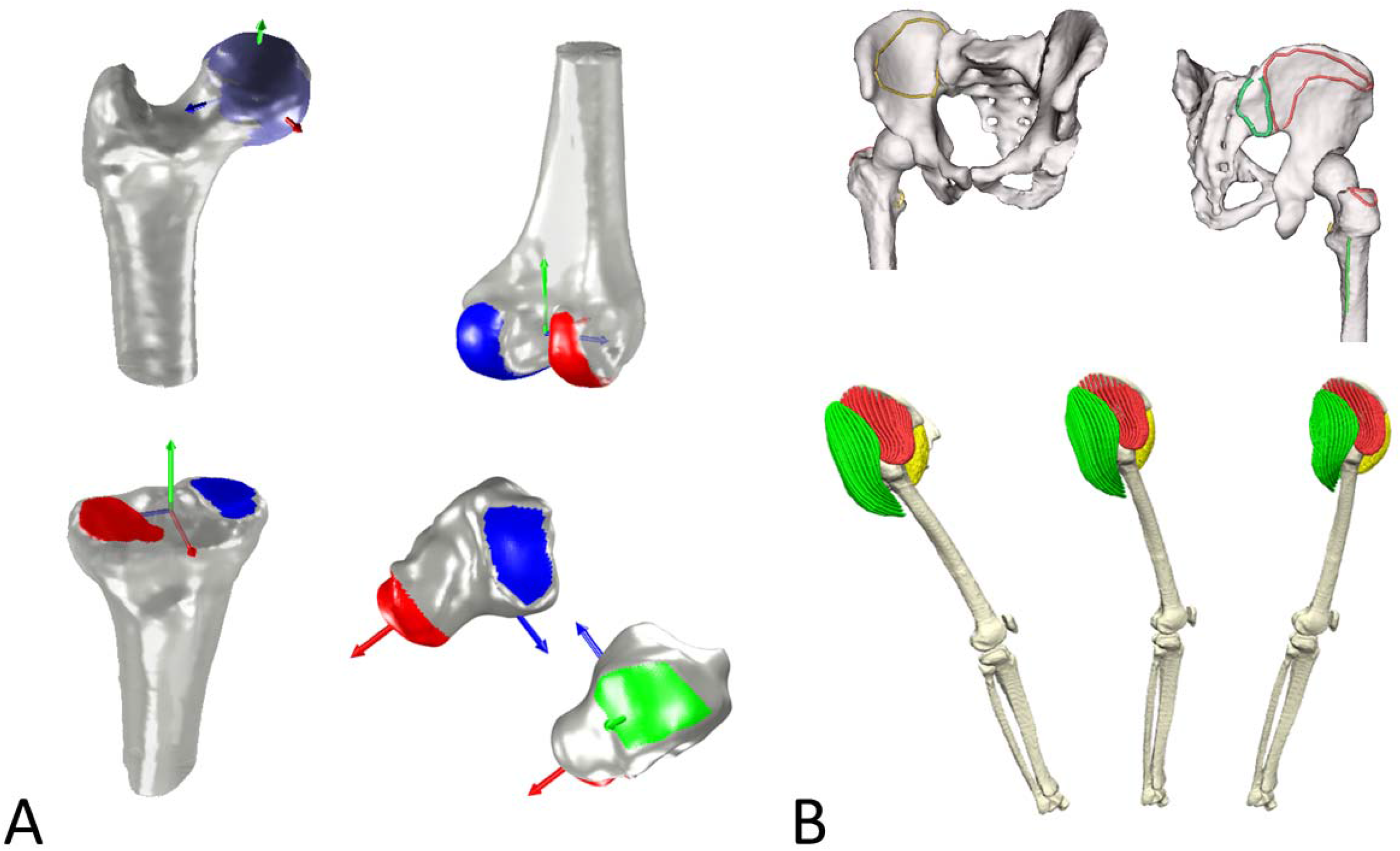
Examples of articular surfaces identified at the femur, tibia and talus by the automatic algorithms (A) and example of augmentation of the ICL-MRI lower extremity model using an automatically generated subject-specific hip musculature including muscle fibres and attachment areas (B).

Extending skeletal models with models of muscle anatomy can also be completely automated (Modenese and Kohout, 2020). In a previous contribution (Modenese et al., 2020) we have used a non-rigid iterative closest point registration (Audenaert et al., 2019) to map the muscle attachment areas from a cadaveric dataset to the ICL-MRI participant’s bones, generating highly-discretized, personalised muscle representations from segmented muscle geometries and simulating their kinematics during gait (Figure 6-B). Future efforts will focus on streamlining these methodologies towards a comprehensive, fully automated, modelling tool for generating subject-specific musculoskeletal models.

In summary, this work presents a computational tool enabling researchers to generate articulated, subject-specific skeletal models of the lower limb in negligible time through a completely automatic workflow that takes three-dimensional bone geometries as inputs. These models can be used immediately for kinematic and kinetic analyses or can serve as extendable baselines for complete musculoskeletal models including musculotendon actuators. This work is framed in a long-term plan aiming to advance the state of the art of anatomical modelling and promote large-scale clinical adoption of personalised computational models of the musculoskeletal system through complete automation of the most challenging modelling tasks.

All data, models and scripts used in this paper are available for download, as detailed in the Appendix.

## CRediT authorship contribution statement

### Luca Modenese

Conceptualization, Methodology, Software, Data Curation, Validation, Formal analysis, Writing – original draft, Writing – review and editing.

### Jean-Baptiste Renault

Investigation, Software, Writing – review and editing.

## Acknowledgements

LM was supported by an Imperial College Research Fellowship granted by Imperial College London and by an Academy of Medical Sciences Springboard Grant [SBF004\1056] supported by the Academy of Medical Sciences, the British Heart Foundation, Diabetes UK, the Global Challenges Research Fund, the Government Department for Business, Energy and Industrial Strategy and the Wellcome Trust. The authors want to thank Dr Michael Rainbow for making available the MATLAB code employed in the Miranda-Femur and Miranda-Tibia algorithms. LM wants to thank Metin Bicer, Arnault Caillet, Bobby Zhang, Clement Favier and Andrew Phillips for the feedback on the manuscript.

## Appendix

To facilitate the reproducibility and replication of our results, we have released our research code and data with this publication. STAPLE is openly developed and shared under the Apache License 2.0 via its repository at https://github.com/modenaxe/msk-STAPLE. All of the data and scripts needed to run the calculations reported in this work, as well as the post-processing scripts to reproduce the figures in this paper and the conference abstracts of the authors mentioned in the main text are available at https://github.com/modenaxe/auto-lowerlimb-models-paper. The version of the scripts and data used in this paper are available through Zenodo at [link will be added upon acceptance] and at https://simtk.org/projects/auto-sk-models.

<u>**NOTE: Public links will be activated upon paper acceptance.**</u>

## References

Audenaert, E.A., Van Houcke, J., Almeida, D.F., Paelinck, L., Peiffer, M., Steenackers, G., Vandermeulen, D., 2019. Cascaded statistical shape model based segmentation of the full lower limb in CT. Computer Methods in Biomechanics and Biomedical Engineering 22, 644–657.

Barber, L., Carty, C., Modenese, L., Walsh, J., Boyd, R., Lichtwark, G., 2017. Medial gastrocnemius and soleus muscle◻tendon unit, fascicle, and tendon interaction during walking in children with cerebral palsy. Developmental Medicine & Child Neurology 59, 843–851.

Barzan, M., Modenese, L., Carty, C.P., Maine, S., Stockton, C.A., Sancisi, N., Lewis, A., Grant, J., Lloyd, D.G., da Luz, S.B., 2019. Development and validation of subject-specific pediatric multibody knee kinematic models with ligamentous constraints. Journal of biomechanics 93, 194–203.

Brandon, S.C., Smith, C.R., Thelen, D.G., 2017. Simulation of soft tissue loading from observed movement dynamics, Handbook of Human Motion. Springer, pp. 1–34.

Brito da Luz, S., Modenese, L., Sancisi, N., Mills, P.M., Kennedy, B., Beck, B.R., Lloyd, D.G., 2017. Feasibility of using MRIs to create subject-specific parallel-mechanism joint models. Journal of Biomechanics 53, 45–55.

Clarke, S., Cobb, J., Jaere, M., Jones, G., Kley, K., Lobenhoffer, P., McCrum, C., Musahl, V., Takeuchi, R., 2018. Osteotomies: Advanced and Complex Techniques, ESSKA Instructional Course Lecture Book. Springer, pp. 129–151.

Conconi, M., Leardini, A., Parenti-Castelli, V., 2015. Joint kinematics from functional adaptation: A validation on the tibio-talar articulation. Journal of Biomechanics 48, 2960–2967.

Damsgaard, M., Rasmussen, J., Christensen, S.T., Surma, E., de Zee, M., 2006. Analysis of musculoskeletal systems in the AnyBody Modeling System. Simulation Modelling Practice and Theory 14, 1100–1111.

Davico, G., Pizzolato, C., Killen, B.A., Barzan, M., Suwarganda, E.K., Lloyd, D.G., Carty, C.P., 2019. Best methods and data to reconstruct paediatric lower limb bones for musculoskeletal modelling. Biomechanics and Modeling in Mechanobiology.

Delp, S.L., Anderson, F.C., Arnold, A.S., Loan, P., Habib, A., John, C.T., Guendelman, E., Thelen, D.G., 2007. OpenSim: open-source software to create and analyze dynamic simulations of movement. IEEE Transactions on Biomedical Engineering 54, 1940–1950.

Dembia, C.L., Bianco, N.A., Falisse, A., Hicks, J.L., Delp, S.L., 2019. OpenSim Moco: Musculoskeletal optimal control. BioRxiv, 839381.

Falisse, A., Pitto, L., Kainz, H., Hoang, H., Wesseling, M., Van Rossom, S., Papageorgiou, E., Bar-On, L., Hallemans, A., Desloovere, K., Molenaers, G., Van Campenhout, A., De Groote, F., Jonkers, I., 2020. Physics-Based Simulations to Predict the Differential Effects of Motor Control and Musculoskeletal Deficits on Gait Dysfunction in Cerebral Palsy: A Retrospective Case Study. Frontiers in Human Neuroscience 14.

Falisse, A., Serrancolí, G., Dembia, C.L., Gillis, J., Jonkers, I., De Groote, F., 2019. Rapid predictive simulations with complex musculoskeletal models suggest that diverse healthy and pathological human gaits can emerge from similar control strategies. Journal of The Royal Society Interface 16, 20190402.

Fischer, M.C.M., Krooß, F., Habor, J., Radermacher, K., 2019. A robust method for automatic identification of landmarks on surface models of the pelvis. Scientific Reports 9, 13322.

Fox, A.S., Carty, C.P., Modenese, L., Barber, L.A., Lichtwark, G.A., 2018. Simulating the effect of muscle weakness and contracture on neuromuscular control of normal gait in children. Gait & posture 61, 169–175.

Gonzalez-Ochoa, C., McCammon, S., Peters, J., 1998. Computing moments of objects enclosed by piecewise polynomial surfaces. ACM Transactions on Graphics 17, 143–157.

Hamner, S.R., Seth, A., Delp, S.L., 2010. Muscle contributions to propulsion and support during running. Journal of Biomechanics 43, 2709–2716.

Hannah, I., Montefiori, E., Modenese, L., Prinold, J., Viceconti, M., Mazzà, C., 2017. Sensitivity of a juvenile subject-specific musculoskeletal model of the ankle joint to the variability of operator-dependent input. Proceedings of the Institution of Mechanical Engineers, Part H: Journal of Engineering in Medicine 231, 415–422.

Henckel, J., Richards, R., Lozhkin, K., Harris, S., Baena, F.R.y., Barrett, A., Cobb, J., 2006. Very low-dose computed tomography for planning and outcome measurement in knee replacement. The Journal of bone and joint surgery. British volume 88, 1513–1518.

Hirschmann, A., Buck, F.M., Fucentese, S.F., Pfirrmann, C.W., 2015. Upright CT of the knee: the effect of weight-bearing on joint alignment. European radiology 25, 3398–3404.

Kai, S., Sato, T., Koga, Y., Omori, G., Kobayashi, K., Sakamoto, M., Tanabe, Y., 2014. Automatic construction of an anatomical coordinate system for three-dimensional bone models of the lower extremities – Pelvis, femur, and tibia. Journal of Biomechanics 47, 1229–1233.

Kainz, H., Modenese, L., Lloyd, D.G., Maine, S., Walsh, H.P.J., Carty, C.P., 2016. Joint kinematic calculation based on clinical direct kinematic versus inverse kinematic gait models. Journal of Biomechanics 49, 1658–1669.

Marra, M.A., Vanheule, V., Fluit, R., Koopman, B.H., Rasmussen, J., Verdonschot, N., 2015. A Subject-Specific Musculoskeletal Modeling Framework to Predict in Vivo Mechanics of Total Knee Arthroplasty. Journal of Biomechanical Engineering 137, 020904.

Martelli, S., Valente, G., Viceconti, M., Taddei, F., 2014. Sensitivity of a subject-specific musculoskeletal model to the uncertainties on the joint axes location. Computer Methods in Biomechanics and Biomedical Engineering, 1–9.

Matsiushevich, K., Belvedere, C., Leardini, A., Durante, S., 2019. Quantitative comparison of freeware software for bone mesh from DICOM files. Journal of Biomechanics 84, 247–251.

McConville, J.T., Churchill, T.D., Kaleps, I., Clauser, C.E., Cuzzi, J., 1980. Anthropometric Relationships of Body and Body Segment Moments of Inertia, Tech. Rep. AFAMRL-TR-80-119. Aerospace Medical Research Laboratory, Wright-Patterson Air Force Base, Dayton, OH.

Miranda, D.L., Rainbow, M.J., Leventhal, E.L., Crisco, J.J., Fleming, B.C., 2010. Automatic determination of anatomical coordinate systems for three-dimensional bone models of the isolated human knee. Journal of Biomechanics 43, 1623–1626.

Mirtich, B., 1996. Fast and Accurate Computation of Polyhedral Mass Properties. Journal of Graphics Tools 1, 31–50.

Modenese, L., Caillet, A., Favier, C., Phillips, A., Kohout, J., Year Simulation of the hip muscles kinematics during gait using highly discretised anatomical models. In 26th Congress of the European Society of Biomechanics. Milan 2020 (Rescheduled due to COVID-19 pandemic, abstract available at https://simtk.org/projects/auto-sk-models).

Modenese, L., Ceseracciu, E., Reggiani, M., Lloyd, D.G., 2016. Estimation of musculotendon parameters for scaled and subject specific musculoskeletal models using an optimization technique. Journal of Biomechanics 49, 141–148.

Modenese, L., Jaere, M., Jones, G.G., Phillips, A., McGregor, A.H., Cobb, J.P., 2019. Integration of external knee joint loads in the pre-surgical planning of high tibial osteotomy: a proof-of-concept study XXVII Congress of the International Society of Biomechanics (ISB2019) and 43rd Annual Meeting of the American Society of Biomechanics (ASB2019), abstract available at https://simtk.org/projects/auto-sk-models, Calgary, Canada.

Modenese, L., Kohout, J., 2020. Automated Generation of Three-Dimensional Complex Muscle Geometries for Use in Personalised Musculoskeletal Models. Annals of Biomedical Engineering 48, 1793–1804.

Modenese, L., Montefiori, E., Wang, A., Wesarg, S., Viceconti, M., Mazzà, C., 2018. Investigation of the dependence of joint contact forces on musculotendon parameters using a codified workflow for image-based modelling. Journal of Biomechanics 73, 108–118.

Montefiori, E., Modenese, L., Di Marco, R., Magni-Manzoni, S., Malattia, C., Petrarca, M., Ronchetti, A., de Horatio, L.T., van Dijkhuizen, P., Wang, A., 2019a. An image-based kinematic model of the tibiotalar and subtalar joints and its application to gait analysis in children with Juvenile Idiopathic Arthritis. Journal of biomechanics 85, 27–36.

Montefiori, E., Modenese, L., Di Marco, R., Magni-Manzoni, S., Malattia, C., Petrarca, M., Ronchetti, A., De Horatio, L.T., van Dijkhuizen, P., Wang, A., 2019b. Linking Joint Impairment and Gait Biomechanics in Patients with Juvenile Idiopathic Arthritis. Annals of biomedical engineering 47, 2155–2167.

Nardini, F., Belvedere, C., Sancisi, N., Conconi, M., Leardini, A., Durante, S., Parenti-Castelli, V., 2020. An Anatomical-Based Subject-Specific Model of In-Vivo Knee Joint 3D Kinematics From Medical Imaging. Applied Sciences 10, 2100.

Noguchi, S., Nishio, M., Yakami, M., Nakagomi, K., Togashi, K., 2020. Bone segmentation on whole-body CT using convolutional neural network with novel data augmentation techniques. Computers in Biology and Medicine 121, 103767.

Nolte, D., Ko, S.-T., Bull, A.M.J., Kedgley, A.E., 2020. Reconstruction of the lower limb bones from digitised anatomical landmarks using statistical shape modelling. Gait & Posture 77, 269–275.

Nolte, D., Tsang, C.K., Zhang, K.Y., Ding, Z., Kedgley, A.E., Bull, A.M.J., 2016. Non-linear scaling of a musculoskeletal model of the lower limb using statistical shape models. Journal of Biomechanics 49, 3576–3581.

Pitto, L., Kainz, H., Falisse, A., Wesseling, M., Van Rossom, S., Hoang, H., Papageorgiou, E., Hallemans, A., Desloovere, K., Molenaers, G., Van Campenhout, A., De Groote, F., Jonkers, I., 2019. SimCP: A Simulation Platform to Predict Gait Performance Following Orthopedic Intervention in Children With Cerebral Palsy. Frontiers in Neurorobotics 13.

Rainbow, M.J., Miranda, D.L., Cheung, R.T.H., Schwartz, J.B., Crisco, J.J., Davis, I.S., Fleming, B.C., 2013. Automatic determination of an anatomical coordinate system for a three-dimensional model of the human patella. Journal of Biomechanics 46, 2093–2096.

Renault, J.-B., Aüllo-Rasser, G., Donnez, M., Parratte, S., Chabrand, P., 2018. Articular-surface-based automatic anatomical coordinate systems for the knee bones. Journal of Biomechanics 80, 171–178.

Sancisi, N., Parenti-Castelli, V., 2011. A new kinematic model of the passive motion of the knee inclusive of the patella. Journal of Mechanisms and Robotics 3, 041003.

Saxby, D.J., Modenese, L., Bryant, A.L., Gerus, P., Killen, B., Fortin, K., Wrigley, T.V., Bennell, K.L., Cicuttini, F.M., Lloyd, D.G., 2016. Tibiofemoral contact forces during walking, running and sidestepping. Gait & Posture 49, 78–85.

Scheys, L., Jonkers, I., Loeckx, D., Spaepen, A., Suetens, P., 2006. Automatic identification of muscle insertion sites in MR images using atlas-based, non-rigid registration. Gait & Posture 24, Supplement 2, S71–S72.

Seth, A., Hicks, J.L., Uchida, T.K., Habib, A., Dembia, C.L., Dunne, J.J., Ong, C.F., DeMers, M.S., Rajagopal, A., Millard, M., Hamner, S.R., Arnold, E.M., Yong, J.R., Lakshmikanth, S.K., Sherman, M.A., Ku, J.P., Delp, S.L., 2018. OpenSim: Simulating musculoskeletal dynamics and neuromuscular control to study human and animal movement. PLOS Computational Biology 14, e1006223.

Sholukha, V., Chapman, T., Salvia, P., Moiseev, F., Euran, F., Rooze, M., Jan, S.V.S., 2011. Femur shape prediction by multiple regression based on quadric surface fitting. Journal of Biomechanics 44, 712–718.

Steger, S., Kirschner, M., Wesarg, S., Year Articulated atlas for segmentation of the skeleton from head & neck CT datasets. In 9th IEEE International Symposium on Biomedical Imaging (ISBI). Barcelona, Spain.

Suwarganda, E.K., Diamond, L.E., Lloyd, D.G., Besier, T.F., Zhang, J., Killen, B.A., Savage, T.N., Saxby, D.J., 2019. Minimal medical imaging can accurately reconstruct geometric bone models for musculoskeletal models. PLOS ONE 14, e0205628.

Taddei, F., Martelli, S., Valente, G., Leardini, A., Benedetti, M.G., Manfrini, M., Viceconti, M., 2012. Femoral loads during gait in a patient with massive skeletal reconstruction. Clinical Biomechanics 27, 273–280.

Valente, G., Crimi, G., Vanella, N., Schileo, E., Taddei, F., 2017a. nmsBuilder: Freeware to create subject-specific musculoskeletal models for OpenSim. Computer Methods and Programs in Biomedicine 152, 85–92.

Valente, G., Pitto, L., Schileo, E., Piroddi, S., Leardini, A., Manfrini, M., Taddei, F., 2017b. Relationship between bone adaptation and in-vivo mechanical stimulus in biological reconstructions after bone tumor: A biomechanical modeling analysis. Clinical Biomechanics 42, 99–107.

Valente, G., Pitto, L., Testi, D., Seth, A., Delp, S.L., Stagni, R., Viceconti, M., Taddei, F., 2014. Are Subject-Specific Musculoskeletal Models Robust to the Uncertainties in Parameter Identification? PLoS ONE 9, e112625.

Victor, J., Premanathan, A., 2013. Virtual 3D planning and patient specific surgical guides for osteotomies around the knee: a feasibility and proof-of-concept study. The bone & joint journal 95, 153–158.

Wesseling, M., de Groote, F., Jonkers, I., 2014. The effect of perturbing body segment parameters on calculated joint moments and muscle forces during gait. Journal of Biomechanics 47, 596–601.

Winter, D.A., 2009. Biomechanics and motor control of the human movement, 4 ed. John Wiley & Sons, Waterloo, Ontario, Canada.

Yin, L., Chen, K., Guo, L., Cheng, L., Wang, F., Yang, L., 2015. Identifying the Functional Flexion-extension Axis of the Knee: An In-Vivo Kinematics Study. PloS one 10, e0128877.

Yushkevich, P.A., Piven, J., Hazlett, H.C., Smith, R.G., Ho, S., Gee, J.C., Gerig, G., 2006. User-guided 3D active contour segmentation of anatomical structures: significantly improved efficiency and reliability. Neuroimage 31, 1116–1128.

Zhang, J., Fernandez, J., Hislop-Jambrich, J., Besier, T.F., 2016. Lower limb estimation from sparse landmarks using an articulated shape model. Journal of Biomechanics 49, 3875–3881.

