## supplementary materials for "Automatic Generation of Personalised Skeletal Models of the Lower Limb from Three-Dimensional Bone Geometries"

### Web Supplementary Materials

#### 1 STAPLE algorithms

Since the scripts implementing the automated methods will be made publicly available, the algorithms developed for this study are only summarily described.

**STAPLE-Pelvis**[1]: the convex hull for the pelvis geometry is computed (excluding sacrum) and the largest triangle of the hull identified (Figure S1-A). Its vertices are the anterior superior iliac spines and one of the pubic symphyses. These landmarks are used to identify the cranial, anterior and right directions, and define a reference system with axes aligned with the principal axes of inertia and pointing in those directions. The posterior superior iliac spines are identified as the most posterior points according to those axes. A reference system can now be defined following the recommendation of Wu et al. (2002).

**STAPLE-Talus**: the principal axes of inertia are calculated, and the bone geometry is sliced along its long axis, corresponding to the smallest moment of inertia. The areas of the sliced sections evolve along this axis with a two peaked profile, that when fitted by a bi-gaussian curve, allows to identify the anterior-posterior direction. The following step is projecting the vertices of the triangular mesh on a plane perpendicular to the anterior-posterior axis and enclose them using a quadrilateral polygon. The smallest side of the polygon identifies the cranial aspect of the talus and an axis is defined perpendicular to the anterior-posterior axis and pointing in that direction. The third axis is perpendicular to the other two. In this auxiliary reference system, the approximate locations of the talocrural and subtalar articular surfaces are known. The talar trochlea articular surface is then identified by analyzing the change of curvature, which is maximum at the articular surface edges, and a cylinder is fitted to the vertices of this area. The axis of the cylinder is the talocrural axis (or ankle axis). The talonavicular surface is identified by analyzing the normals of the elements of the most anterior quarter of the talus, while the talocalcaneal surface with an analysis of curvature of the posterior part of the talus. A sphere is fitted to each of the identified surfaces and the subtalar axis is identified as the line connecting the centers of the two spheres.

**STAPLE-Foot**: using all the foot bones except the phalanges, a convex hull is computed and similarly to the pelvis, the largest triangle is identified. The vertices of this triangles are roughly the points of contact of the calcaneus and first and fifth metatarsal bones with the ground. These points are used to construct an auxiliary reference system with an X axis pointing anteriorly from the heel to the midpoint of the two anterior points and a Z axis lying on the same triangle but perpendicular to X. These axes are used to identify the markers HEE (most posterior point) and D1M and D5M landmarks (most medial and most lateral points).

---

[1] We discovered after completing our manuscript that a similar algorithm had been independently proposed by Seim et al. (2009) in a symposium proceeding.

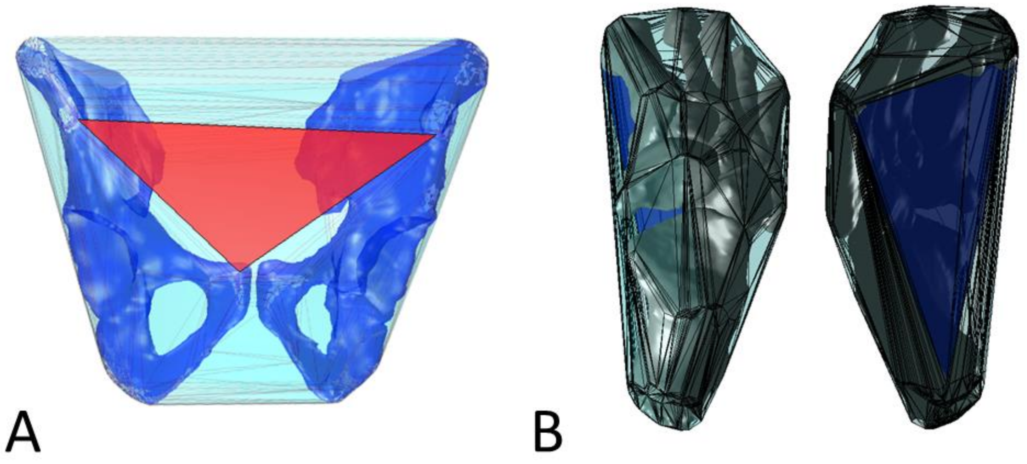


*Figure S1 Convex hull used to build the reference system of the pelvis in the STAPLE-Pelvis algorithm and largest triangle (in red) used for constructing the pelvis-ground coordinate system (A). Convex hull used to identify the foot sole in the STAPLE-Foot algorithm and largest triangle (in blue) used to define the foot sole reference system (B), seen from above (left) and from below (right).*

### 2 Generation of partial models

The STAPLE toolbox can generate partial models of the lower limb, depending on the availability of bone geometries. Figure S2 demonstrates this functionality using the datasets employed in the main manuscript and the ankle and foot bone geometries shared by Montefiori et al (2019a) and available in folder “Data\Repeatability Study\Fitting3x3\P2\Geometries” of the public package of supplementary materials downloadable at *https://doi.org/10.15131/shef.data.5863443.v1*. These partial models and the scripts employed to generate them are available in the STAPLE Github repository (https://github.com/modenaxe/msk-STAPLE) as examples of use of the toolbox.

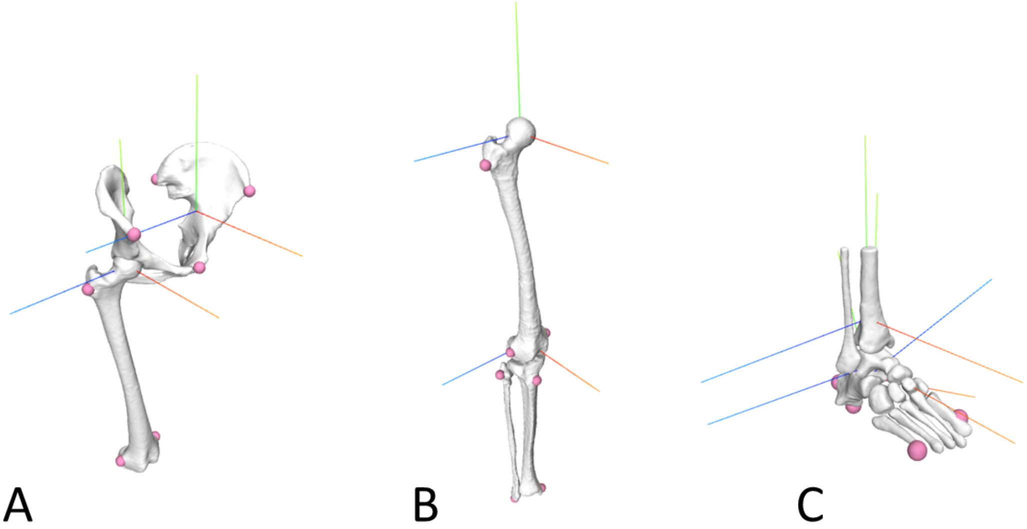


*Figure S2 Example of partial models generated using STAPLE: a hip model (A), a tibiofemoral model (B) and a model of the ankle complex (C). The last model was generated using the data released as Supplementary materials by Montefiori et al. (2019). The child reference systems are displayed for all joints included in the models.*

### 3 Estimation of femoral and tibial longitudinal axis using principal component analysis versus principal axes of inertia

In Table S1 we present differences in the estimation of the anatomical vertical axis of femur and tibia when using a principal component (PC) approach (as in the Kai-algorithms) and a principal axis of inertia (PIA) (as in the GIBOC-algorithms). This comparison is of interest in the femur and tibia because the vertical axis is employed in the GIBOC-algorithms as auxiliary vector for the definition of joint coordinate systems (JCSs), while in the Kai-algorithms it represents the cranial-caudal Y axis of the JCSs. Therefore, we wanted to verify if the two approaches yield substantially different results and how the difference varied depending on the level of completeness of the bone geometries (full bones, both epiphyses without diaphysis and distal femur and proximal tibia only, as shown in Figure S3). The results will contribute to explain the difference on Y-axis reported in Table 5 of the manuscript. Table S1 shows differences <3° and <5° between the two approaches when the complete femurs and tibias are considered respectively. Interestingly the vertical axis estimations of PCA and PIA become more similar to each other when the diaphysis is removed from the femoral geometry, or the distal femur is considered. The opposite trend is observed for the tibia, with less compatible estimations for incomplete bone geometries.

These results suggest caution in comparing simulation results obtained from STAPLE models including JCSs computed from complete bone geometries with those from STAPLE models built from partial bone geometries because there could be systematic offsets in the estimations of the longitudinal axes of the long bones.

Table S1 Estimation of the anatomical long axis of long bones when using principal component analysis or principal axis of inertia with different levels of bone geometry completeness.

| | **Dataset** | **PCA versus PIA difference [deg]** | | |
|---|---|---|---|---|
| | | entire bones | distal femur/ proximal tibia | bone epiphyses |
| femur | LHDL-CT | 2.9 | 1.5 | 2.2 |
| | TLEM2-CT | 1.6 | 1.6 | 0.8 |
| | ICL-MRI | 2.9 | 0.9 | 2.1 |
| | JIA-MRI | 2.8 | 1.1 | 2.4 |
| tibia | LHDL-CT | 4.9 | 10.6 | 6.7 |
| | TLEM2-CT | 0.3 | 6.5 | 2.6 |
| | ICL-MRI | 3.6 | 8.3 | 5.2 |
| | JIA-MRI | 3.9 | 11.9 | 4.8 |

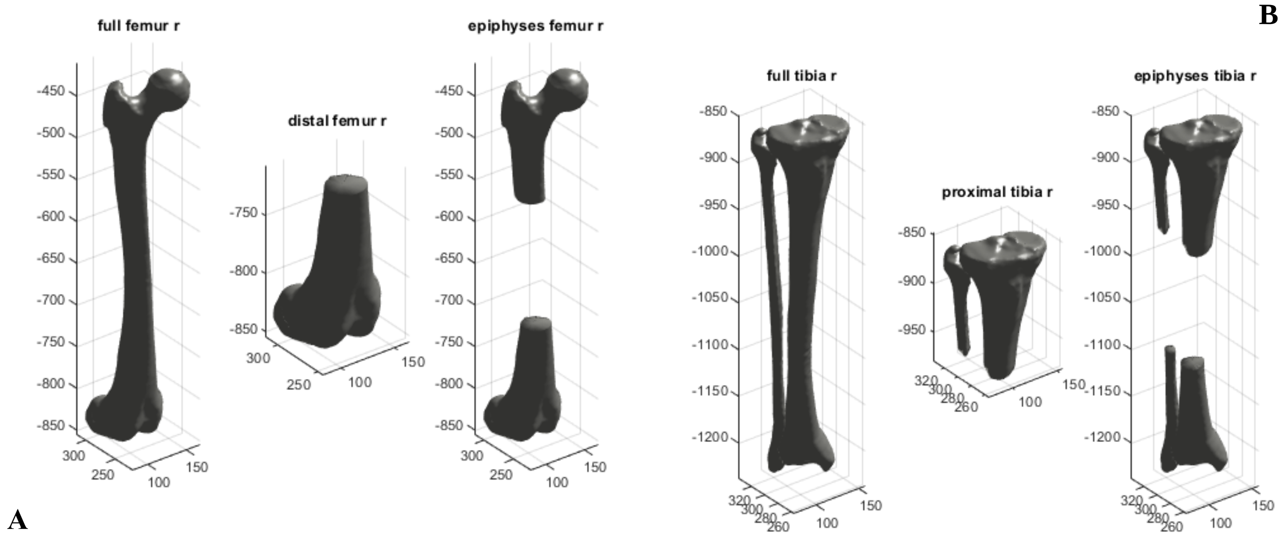


*Figure S3 Bone geometries of femur (A) and tibia (B) with various level of completeness, as considered in Table S2: full bones (left), just one epiphysis (center) or long bones without diaphysis (right).*

### 4 Detailed results from gait simulations

This section presents the correlation coefficients, root mean squared errors (RMSEs), and the results of the statistical parametric mapping (SPM) two-tailed t-test for the joint angles (Table S2) and joint moments (Table S3) curves obtained from the gait simulations described in the manuscript. Tables S2-S3 complement the results reported in the article by presenting the metric values for each of the coordinates of the considered models. Finally, Figures S4-S8 present the t-statistics calculated by the SPM t-test and display the areas of the gait cycle where significant differences were identified.

Table S2 Mean values (and standard deviations) of the correlation coefficients and RMSEs for all the joint angles of the automatic and manual JIA-MRI models calculated across the six available walking simulations, and total length of the clusters in which the two curve sets are significantly different according to the SPM t-test, expressed as percentage of the gait cycle (last column).

| **Joint Angle** | **Correlation coefficient* (mean)** | **Correlation coefficient (std)** | **RMSE (mean) [deg]** | **RMSE (std) [deg]** | **SPM t-test cluster length [% gait cycle]** |
|---|---|---|---|---|---|
| Pelvic tilt | 0.997 | 0.001 | 3.4 | 0.01 | 100 |
| Pelvic rotation | 0.996 | 0.001 | 1.0 | 0.03 | 32 |
| Pelvic list | 0.999 | 0.000 | 0.2 | 0.02 | 0 |
| Hip flex/extension | 1.000 | 0.000 | 3.4 | 0.01 | 96 |
| Hip ad/abduction | 1.000 | 0.000 | 1.0 | 0.01 | 13 |
| Hip int/external rotation | 0.998 | 0.000 | 0.4 | 0.07 | 0 |
| Knee flex/extension | 1.000 | 0.000 | 0.1 | 0.01 | 0 |
| Ankle dorsi/plantarflexion | 0.998 | 0.000 | 1.3 | 0.04 | 0 |
| Subtalar ev/inversion | 0.998 | 0.000 | 1.6 | 0.11 | 7 |

* All correlation coefficients have $p<<0.0001$

Table S3 Mean values (and standard deviations) of the correlation coefficients and RMSE for all the net joint moments (normalized by body weight) of the automatic and manual JIA-MRI models calculated across the six walking simulations, and total length of the clusters in which the two curve sets are significantly different according to the SPM t-test, expressed as percentage of the gait cycle (last column).

| **Joint moment** | **Correlation coefficient* (mean)** | **Correlation coefficient (std)** | **RMSE (mean) [Nm/Kg]** | **RMSE (std) [Nm/Kg]** | **SPM t-test cluster length [% gait cycle]** |
|---|---|---|---|---|---|
| Hip flex/extension | 0.973 | 0.001 | 0.078 | 0.003 | 31 |
| Hip ad/abduction | 0.999 | 0.000 | 0.035 | 0.002 | 11 |
| Hip int/external rotation | 0.934 | 0.009 | 0.028 | 0.002 | 27 |
| Knee flex/extension | 0.994 | 0.001 | 0.033 | 0.001 | 29 |
| Ankle dorsi/plantarflexion | 1.000 | 0.000 | 0.007 | 0.000 | 37 |
| Subtalar ev/inversion | 0.996 | 0.001 | 0.012 | 0.001 | 11 |

* All correlation coefficients have p<<0.0001

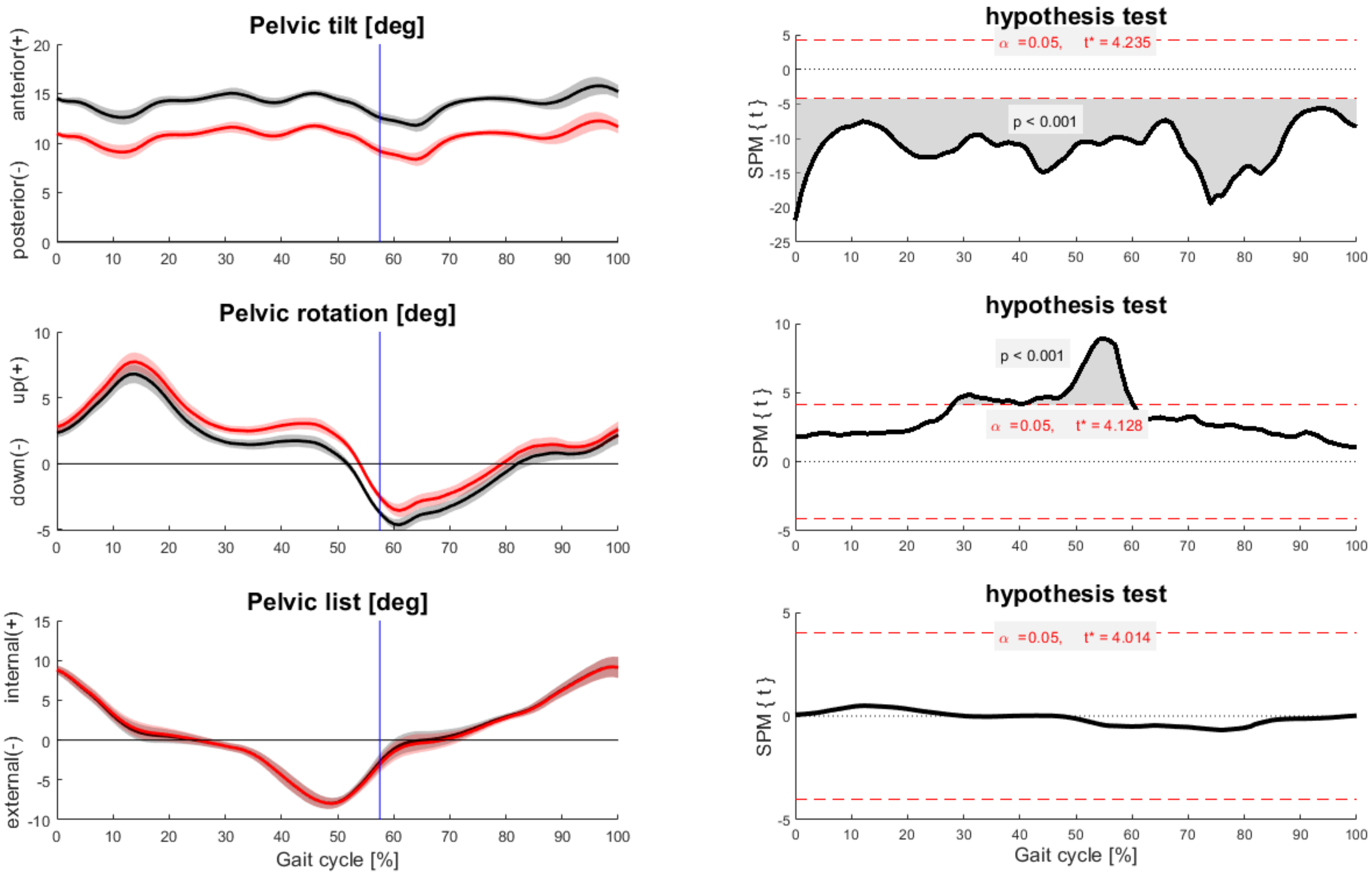


*Figure S4 Joint angles for the pelvis-ground joint obtained from six walking simulations performed with the automatic (red lines) and manual (black lines) JIA-MRI models (left column) and correspondent t statistics calculated using the statistical parametric mapping equivalent of a paired, two-tailed t-test with significance level set to α=0.05 (right column). Grey shaded areas identify regions of the gait cycle where significant differences exist between the joint angles (p values reported in proximity of the clusters).*

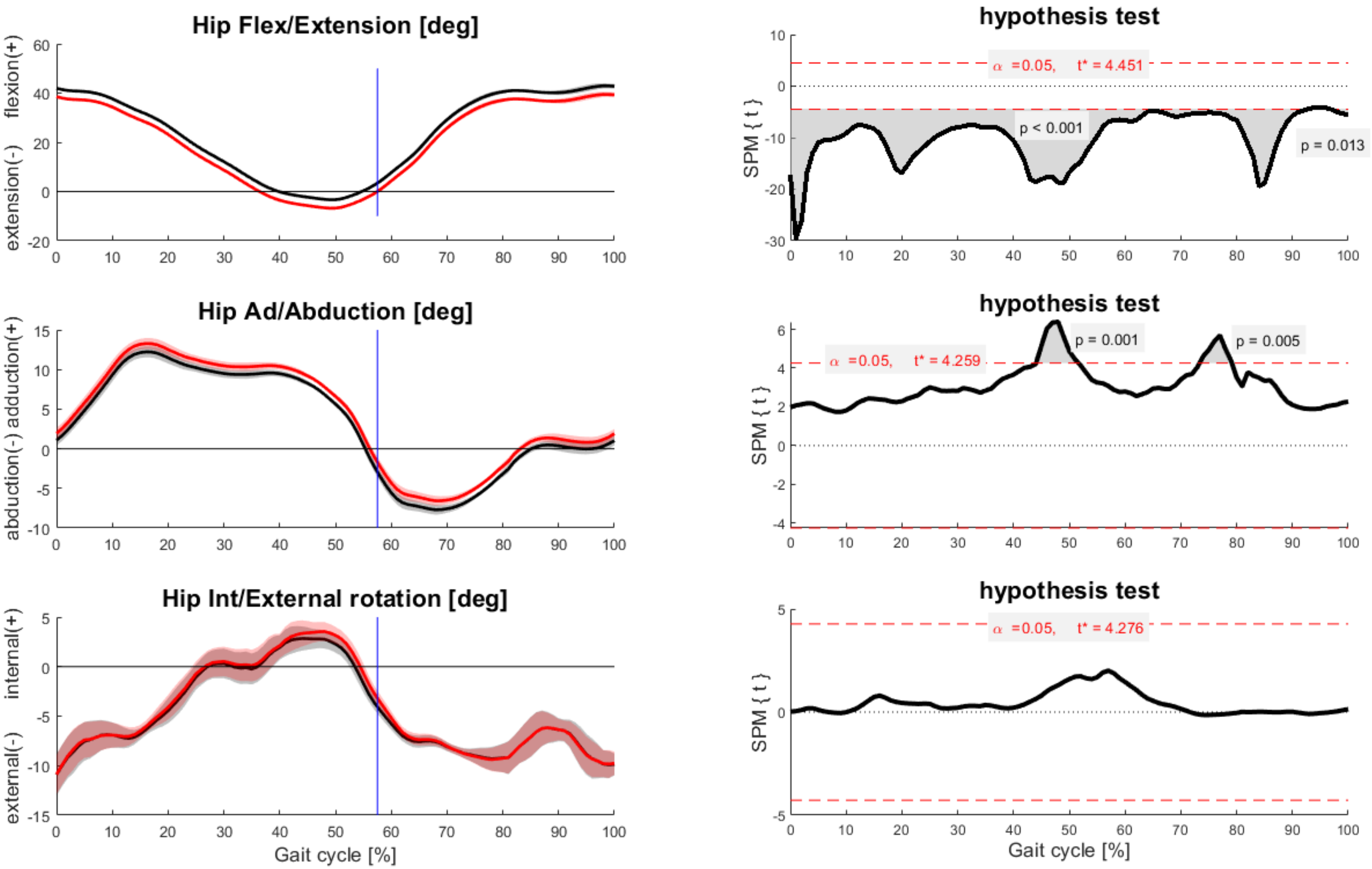


*Figure S5 Joint angles for the hip joint obtained from six walking simulations performed with the automatic (red lines) and manual (black lines) JIA-MRI models (left column) and correspondent t statistics calculated using the statistical parametric mapping equivalent of a paired, two-tailed t-test with significance level set to α=0.05 (right column). Grey shaded areas identify regions of the gait cycle where significant differences exist between the joint angles (p values reported in proximity of the clusters).*

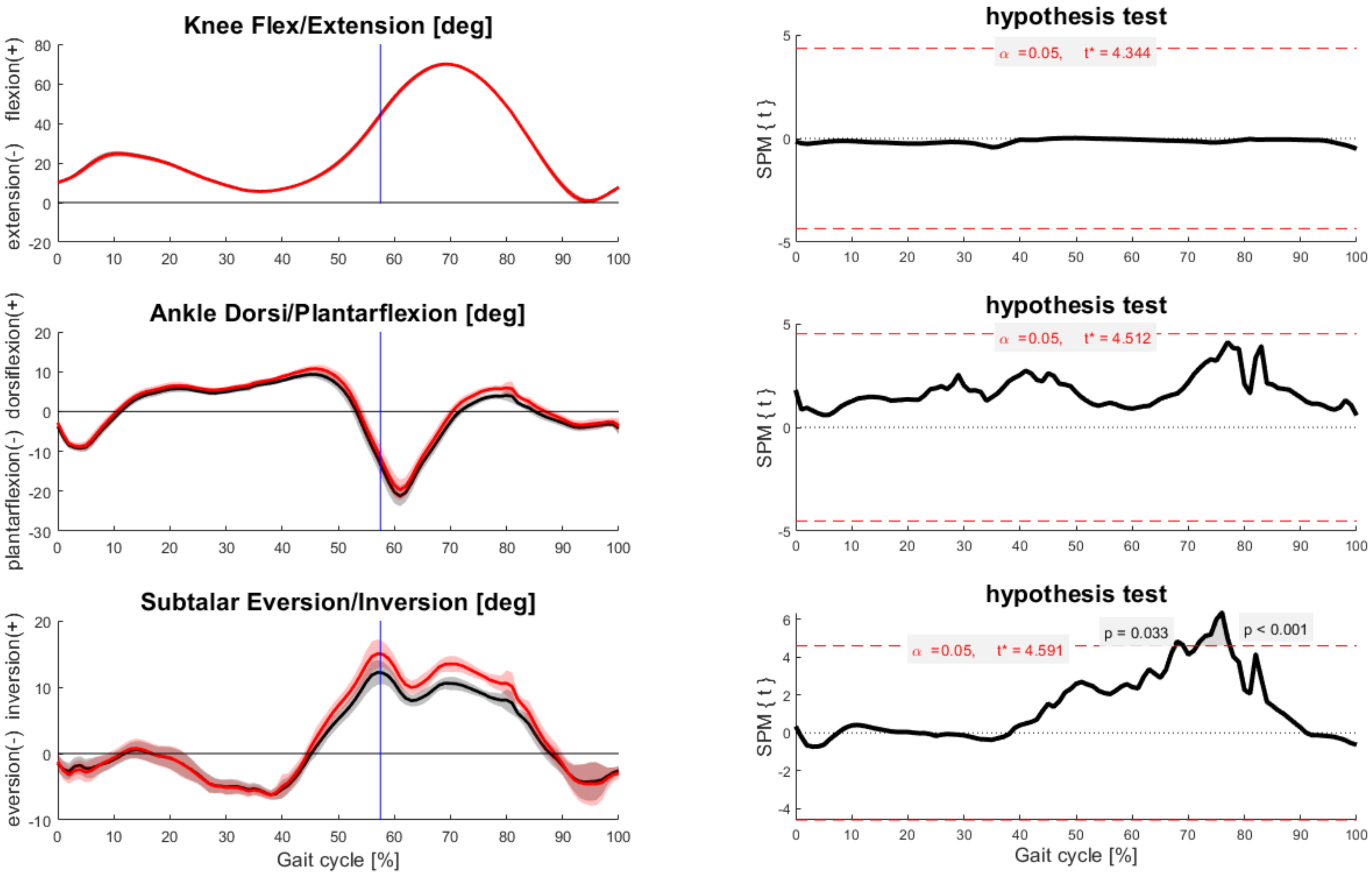


*Figure S6 Joint angles for the knee, talocrural and subtalar joints, obtained from six walking simulations performed with the automatic (red lines) and manual (black lines) JIA-MRI models (left column) and correspondent t statistics calculated using the statistical parametric mapping equivalent of a paired, two-tailed t-test with significance level set to α=0.05 (right column). Grey shaded areas identify regions of the gait cycle where significant differences exist between the joint angles (p values reported in proximity of the clusters).*

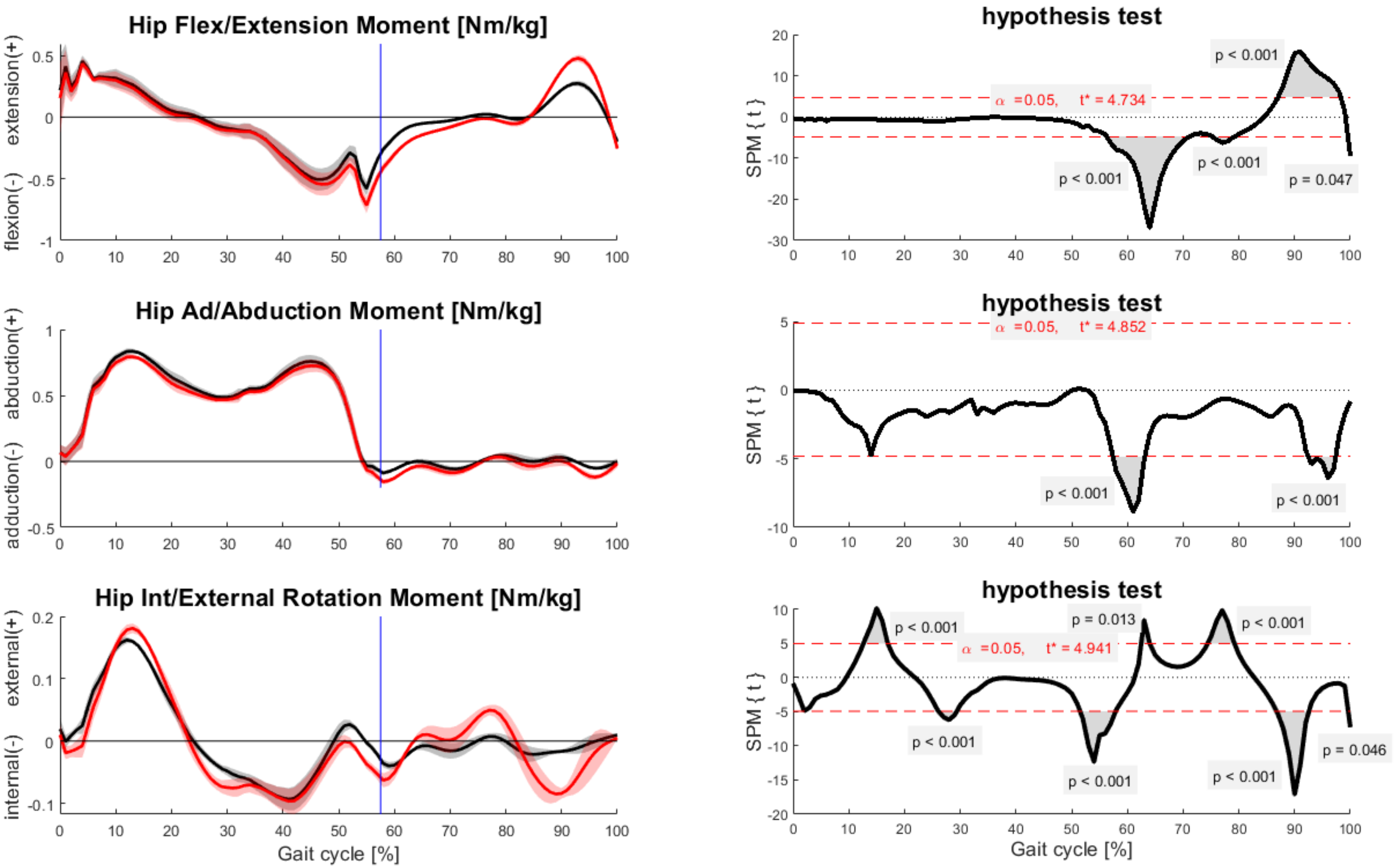


*Figure S7 Joint moments for the hip joint, normalized by body weight, obtained from six walking simulations performed with the automatic (red lines) and manual (black lines) JIA-MRI models (left column) and correspondent t statistics calculated using the statistical parametric mapping equivalent of a paired, two-tailed t-test with significance level set to α=0.05 (right column). Grey shaded areas identify regions of the gait cycle where significant differences exist between the joint angles (p values reported in proximity of the clusters).*

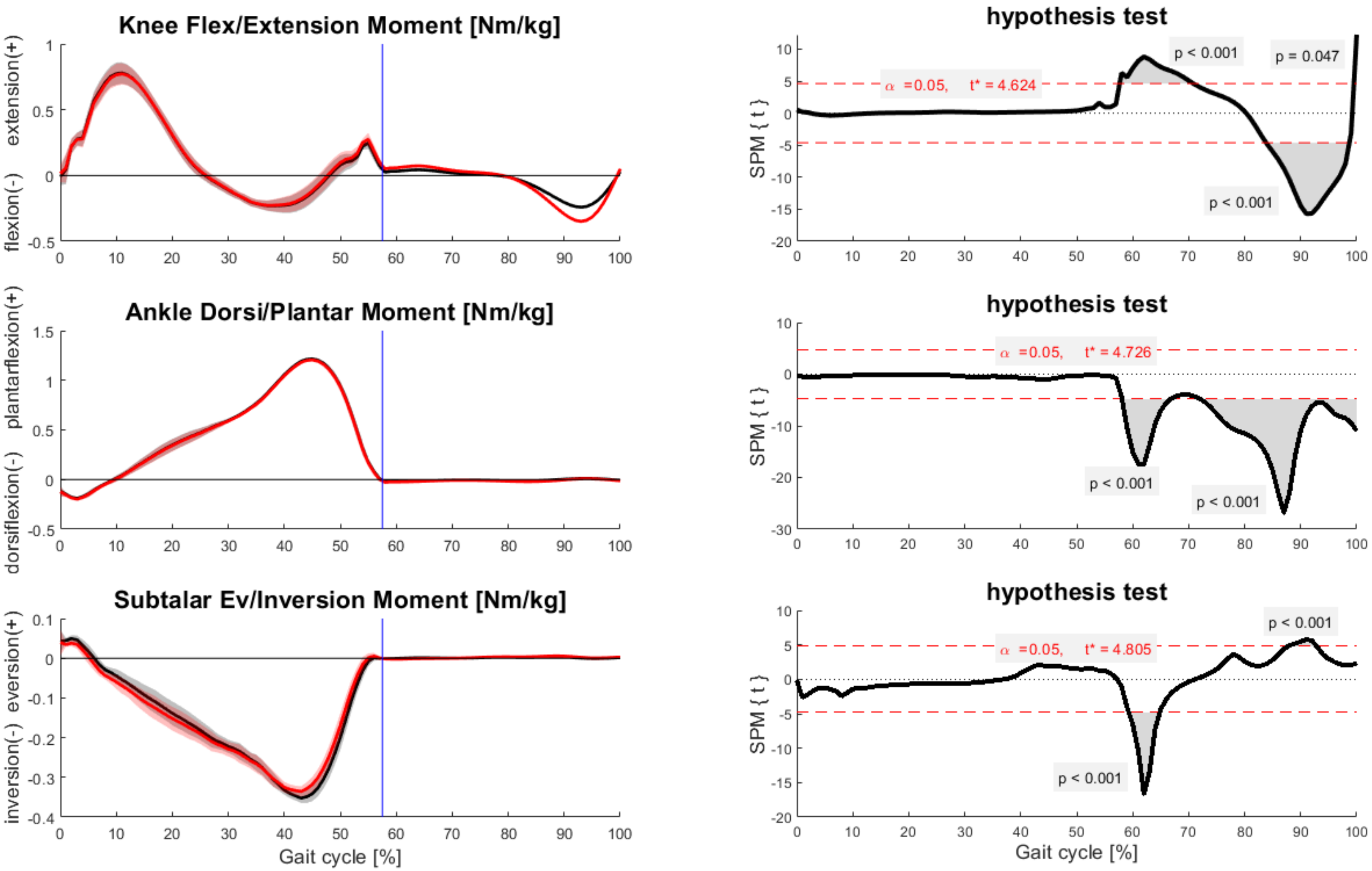


*Figure S8 Joint moments for the knee, talocrural and subtalar joints, normalized by body weight, obtained from six walking simulations performed with the automatic (red lines) and manual (black lines) JIA-MRI models (left column) and correspondent t statistics calculated using the statistical parametric mapping equivalent of a paired, two-tailed t-test with significance level set to α=0.05 (right column). Grey shaded areas identify regions of the gait cycle where significant differences exist between the joint angles (p values reported in proximity of the clusters).*

### REFERENCES

Montefiori, E., Modenese, L., Di Marco, R., Magni-Manzoni, S., Malattia, C., Petrarca, M., Ronchetti, A., de Horatio, L.T., van Dijkhuizen, P., Wang, A., 2019a. An image-based kinematic model of the tibiotalar and subtalar joints and its application to gait analysis in children with Juvenile Idiopathic Arthritis. Journal of Biomechanics 85, 27-36.

Montefiori, E., Modenese, L., Di Marco, R., Magni-Manzoni, S., Malattia, C., Petrarca, M., Ronchetti, A., De Horatio, L.T., van Dijkhuizen, P., Wang, A., 2019b. Linking Joint Impairment and Gait Biomechanics in Patients with Juvenile Idiopathic Arthritis. Annals of biomedical engineering 47, 2155-2167.

Seim, H., Kainmueller, D., Heller, M., Zachow, S., Hege, H.-C., Year Automatic extraction of anatomical landmarks from medical image data: An evaluation of different methods. In 2009 IEEE International Symposium on Biomedical Imaging: From Nano to Macro.
